# Female specific dysfunction of sensory neocortical circuits in a mouse model of autism mediated by mGluR5 and Estrogen Receptor α

**DOI:** 10.1101/2023.08.10.552857

**Authors:** Gemma Molinaro, Jacob E. Bowles, Katilynne Croom, Darya Gonzalez, Saba Mirjafary, Shari Birnbaum, Khaleel A. Razak, Jay R. Gibson, Kimberly M. Huber

**Author notes:** Correspondence (K.M.H.).

## Abstract

Autism manifests differently in males and females and the brain mechanisms that mediate these sex-dependent differences are unknown. Here, we demonstrate that deletion of the ASD-risk gene, *Pten,* in neocortical pyramidal neurons (NSE*Pten* KO) results in robust hyperexcitability of local neocortical circuits in female, but not male, mice, observed as prolonged, spontaneous persistent activity states (UP states). Circuit hyperexcitability in NSE*Pten* KO mice is mediated by enhanced and/or altered signaling of metabotropic glutamate receptor 5 (mGluR5) and estrogen receptor α (ERα) to ERK and protein synthesis selectively in *Pten* deleted female neurons. In support of this idea, *Pten* deleted Layer 5 cortical neurons have female-specific increases in mGluR5 and mGluR5-driven protein synthesis. In addition, mGluR5-ERα complexes are elevated in female cortex and genetic reduction of ERα in *Pten* KO cortical neurons rescues circuit excitability, protein synthesis and enlarged neurons selectively in females. Abnormal timing and hyperexcitability of neocortical circuits in female NSE*Pten* KO mice are associated with deficits in temporal processing of sensory stimuli and social behaviors as well as mGluR5-dependent seizures. Female-specific cortical hyperexcitability and mGluR5-dependent seizures are also observed in a human disease relevant mouse model, germline *Pten*+/- mice. Our results reveal molecular mechanisms by which sex and a high impact ASD-risk gene interact to affect brain function and behavior.

## Introduction

Autism Spectrum Disorder (ASD) is diagnosed 4 times more often in males than females ^1^. However, increasing evidence indicates that ASD is expressed differently in males and females on a behavioral level and affects brain function and connectivity differently. Because of these sex differences, it has been suggested that ASD is not as well detected in females with current ASD diagnostic tests and is more prevalent in females than previously recognized ^2–7^. Some symptoms and co-morbidities, such as deficits in sensory processing and epilepsy, are more severe or common in females with autism ^8–11^. There is little known of the sex-dependent brain mechanisms that interact with ASD-linked genes and how ASD genes affect female brain physiology and behaviors differently than in males. One prominent ASD gene associated with sex-dependent interactions, at least in cancer, is the *Phosphatase and tensin homolog deleted on chromosome 10* (*PTEN*), a lipid phosphatase that dephosphorylates PIP3 and inactivates PI3K and its downstream signaling to Akt and mTOR ^12,13^. PTEN was first discovered as a tumor suppressor and subsequently linked to ASD ^14^. Individuals with germline *PTEN* mutations display variable and complex phenotypes including macrocephaly, autism, intellectual disability (ID), and epilepsy that are collectively termed PTEN Hamartoma syndromes (PTHS)^12,15–18^. Importantly, mutations in PTEN or other suppressors of the PI3K/Akt/mTOR pathway are the most common cause of breast cancers; cancers that are ∼ 100X more prevalent in women than men ^19^. This sex bias is because 70-80% of breast cancers express Estrogen receptor α (ERα) and depend on estrogen signaling for growth ^20^. Mutations in *PTEN* or other genes that lead to hyperactivation of Akt/mTOR in breast cancer cells result in enhanced ERα function ^21–23^. Thus, combination therapies that inhibit ERα and PI3K/mTOR are most therapeutic for breast cancer ^23^.

Although there is not a female sex bias in ASD with germline *PTEN* mutations, as there is with breast cancer, the rate of ASD in females with *PTEN* mutations is high (47%) and similar to males (66%)^17^. A mouse model of PTHS, a germline heterozygous deletion of *Pten,* displays female-specific deficits in social behavior ^18,24,25^. Similarly, human relevant *Pten* mutation in mice, leading to a cytoplasmic predominant expression of PTEN (*Pten^m^*^3m4^), results in a male-selective enhancement in sociability ^26^. The sex-dependent effects of PTEN on brain physiology are unknown, nor the molecular mechanisms that lead to sex-dependent behavior deficits with PTEN deletion. ERα is expressed in brain, including cortical neurons, where it has important regulatory functions on different processes such as cognition, anxiety and addiction ^27,28^ Many of the effects of estrogen on neurons and behavior are mediated by ERα and rely on activity of metabotropic glutamate receptor 5, a Gq coupled glutamate receptor and candidate therapeutic target in ASD. If or how PTEN deletion in neurons affects ERα or mGluR5 function or if these signaling pathways contribute to brain phenotypes in PTEN loss of function models is also unknown.

To explore the effects of *Pten* deletion on brain function, we studied an established mouse model of PTEN deletion with ASD-relevant behaviors, ^NSE^*Pten* KO ^29^. ^NSE^*Pten* KO mice have a mosaic deletion of PTEN primary in Layers 3-5 excitatory neurons. To assess overall cortical circuit function, we measured spontaneous persistent activity states, or UP states, in acute slices of the somatosensory cortex. UP states are driven by the activity of local cortical circuits and are an overall readout of the balance of excitation and inhibition. UP states also reflect the state of maturation of cortical circuits ^30^ ^31^. We have shown that UP state duration is prolonged in another mouse model of ASD, the *Fmr1* knockout (KO) model of Fragile X Syndrome (FXS), reflecting a circuit hyperexcitability. Surprisingly, we observed prolonged UP states in female, but not male, ^NSE^*Pten* KO mice in contrast to our previous work in male *Fmr1* KO mice. Prolonged UP states in female ^NSE^*Pten* KO are mediated by enhanced signaling of mGluR5 and ERα and required de novo protein synthesis. We find that *Pten* deletion in L5 neurons results in enhanced mGluR5 levels, mGluR5 and ERα-driven protein synthesis and mGluR5-dependent seizures, only in females, but not males. Notably, genetic reduction of ERα in L5 neurons corrects circuit hyperexcitability, enhanced protein synthesis and cell size, again, selectively in females. The prolonged cortical circuit activity we observe in slices from female ^NSE^*Pten* KO is associated with a female selective deficit in temporal processing of sensory stimuli as measured with electroencephalographic (EEG) recordings *in vivo* as well as deficits in social behavior. Female specific cortical circuit hyperexcitability and mGluR5-dependent seizures are also observed in a mouse model of PTHS. Our results suggest a model where deletion of PTEN results in a female specific enhancement of mGluR5-ERα function that drives cortical hyperexcitability through ERK and *de novo* protein synthesis. These results may suggest distinct therapeutic strategies for males and females with PTEN mutations and highlight the importance of studies of ASD risk genes in both males and female brain physiology.

## Results

### Hyperexcitable sensory neocortical circuits in female mouse model of Pten deletion

To determine the effects of *Pten* deletion on cortical circuit function, we utilized a previously characterized mouse where *Pten* is deleted with *Neuron-specific enolase* (*NSE*) promoter-driven *cre* (^NSE^*Pten* KO). In brain, NSE-Cre is expressed postnatally (∼postnatal day 2) in discrete mature neuronal populations in layers III to V as well as in neurons in the CA3 region and dentate gyrus of the hippocampus ^29^. In cortex, NSE-Cre is most abundantly expressed in sensory areas and the vast majority of neurons are excitatory based on GABA staining >99%, data not shown). Importantly, ^NSE^*Pten* KO mice show ASD-relevant behaviors such as deficits in social behavior, learning and memory and sensory hypersensitivity and thus may be a good model to study ASD-related circuit alterations with PTEN deletion ^29^. To assess sensory cortical circuit function, we performed extracellular multi-unit recordings of spontaneous persistent activity, or UP states, in acute slices of somatosensory barrel neocortex from postnatal day (P) 18-25 male and female mice (Fig. 1A). Mice were either NSE-Cre(+)/*Ptenfl/fl* or NSE-Cre (-) *Ptenfl/fl* or *Ptenfl/+* littermates, indicated in figures as Cre(+) or Cre(-), respectively (Fig. 1B). UP states are a spontaneous, oscillatory (0.5-1 Hz), synchronized firing of neocortical neuron networks driven by recurrent excitatory and inhibitory synaptic circuits and provide a readout of the intact functioning of neocortical circuits ^32,33^. The duration of UP states measured in Layers (L) 4, was increased by 56% in female slices compared to Cre (-) controls (Fig. 1C). In contrast, slices from Cre(+) males had normal UP state duration. There was no difference in UP state duration between male and female Cre(-) mice. Similar results were obtained if recordings were performed in L2/3 or L5 (Fig. 1C). UP states were longer in Cre (+) females, but not males, in comparison to Cre (-) controls and there was a significant interaction of sex and genotype on UP state duration in both L2/3 (2 way ANOVA; F (1, 73) = 5.555; p< 0.05) and in L5 (F (1, 71) = 7.332; p< 0.01). There was no effect of sex or genotype on the amplitude or frequency of UP states in any layer (Supplementary Table 1). Because similar results were obtained in all layers, subsequent experiments were focused in L4.

**Figure 1.**
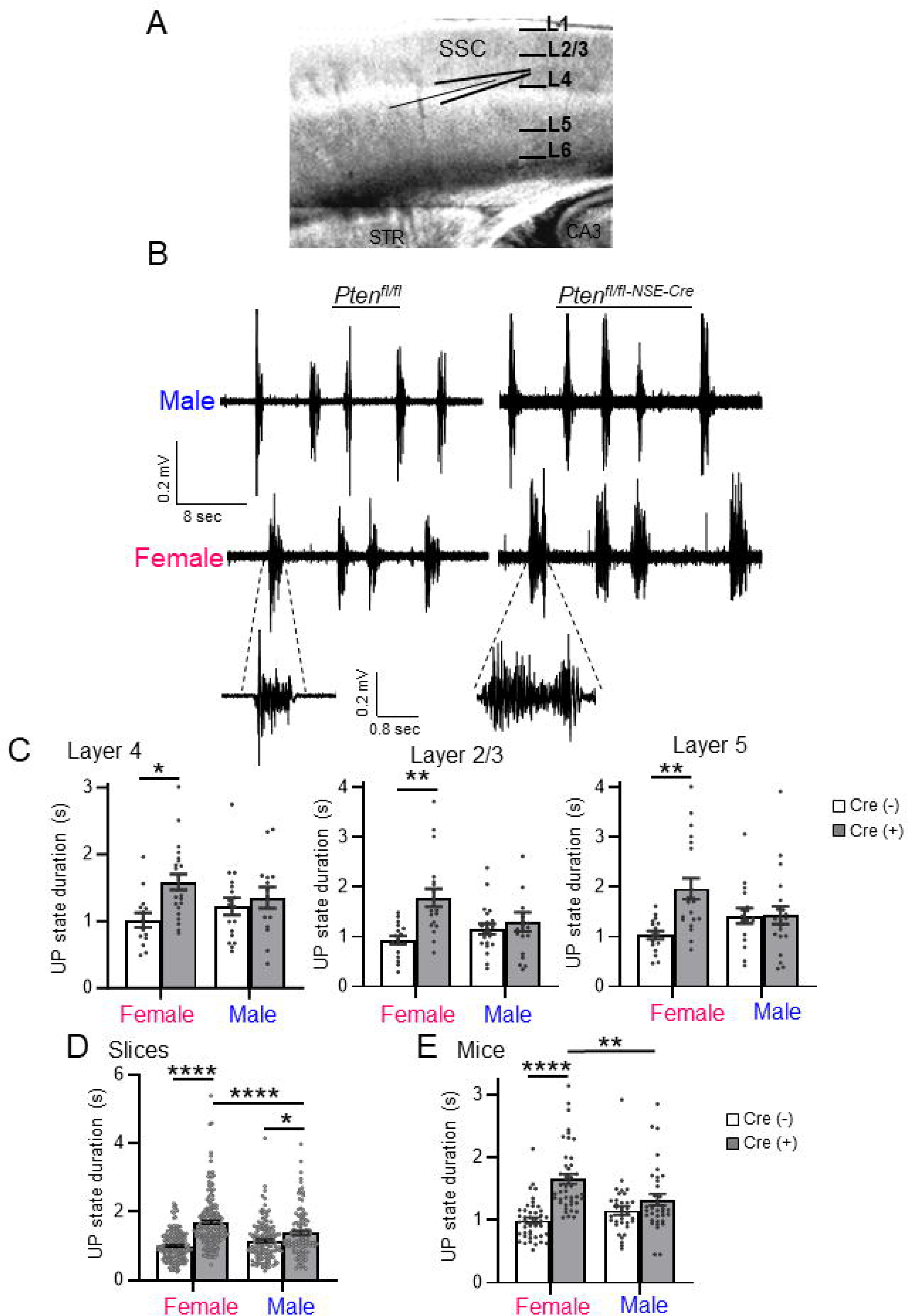
Female specific cortical circuit hyperexcitability in acute slices of ^NSE^*Pten* KO mice. **A.** Schematic of thalamocortical slice of somatosensory cortex and extracellular multiunit recording in Layer (L) 4. **B.** Representative spontaneous, persistent activity states, or UP states from extracellular recordings in L4 of male and female *Pten^fl/fl^* Cre (-) and *Pten^fl/fl-NSE-Cre^*Cre (+) somatosensory cortex. **C.** Group average UP state duration in male and female Cre (-) and Cre (+) mice in L4, L2/3 and L5. (Each black dot is an individual slice from 4-6 mice/sex/genotype). **D, E.** UP state duration reported as average per slice (D) or per mouse (E). (Each black dot is an individual slice (D) or mouse (E) from 35-45 mice/sex/genotype). For all figures, data are represented as mean ± SEM. Two-way ANOVA with Sidak for multiple comparisons. For all figures, ^∗^p < 0.05, ^∗∗^p < 0.01, ^∗∗∗^p < 0.0005, ^∗∗∗∗^p < 0.0001.

In some experiments, we did observe increases in UP state duration in male Cre(+) mice as compared to male Cre(-). Combining data from vehicle or un-treated slices across experiments, we observed a small, ∼20%, genotypic increase in UP state duration in males (Cre(-); 1145±45 ms; n = 148 slices; Cre(+); 1383±62ms; n= 114; p <0.05) (Fig. 1D). However, the genotypic increase in UP state duration in females was large, ∼70%, (Cre (-) 997±31 ms; n = 175; female Cre(+); 1689±58ms; n = 176; p< 0.001) and durations were longer than in Cre(+) males (p< 0.001). No difference in UP state duration was observed between Cre(-) males and females. This resulted in a significant interaction of sex X genotype on UP state duration (F (1, 609) = 20.47; p< 0.0001). Averaging UP state duration from all slices per mouse yielded similar results (Fig. 1E) and a significant sex X genotype interaction (F (1, 153) = 11.58; p< 0.001).

The female specific effects on UP state duration were not attributable to sex-dependent differences in deletion of PTEN in ^NSE^*Pten* KO mice as measured with fluorescent immunohistochemistry or western blotting. Reflecting the expression pattern of NSE-Cre, we observed a mosaic deletion of PTEN in L5 neurons as indicated by NeuN labeling that was similar in males and females (Female Cre(+); 63±2% of NeuN+ L5 neurons; n=14 slices/3 mice; Male Cre(+); 64±2%; n=10 slices/3 mice; Fig. S1A, B). Total PTEN levels were quantified in L5 enriched cortical lysates which revealed similar decreases of PTEN in male and female Cre(+) mice (Female Cre(+); 76±5%; n=10 mice; Male Cre(+); 72±6% of same sex Cre(-); n= 6 mice; Fig. S1C,D). Taken together these results suggest sex specific circuit hyperexcitability in female *Pten*-deleted cortex.

### Pharmacological antagonism of mGluR5 rescues UP states in female Cre(+) mice

Genetic or pharmacological reduction of metabotropic glutamate receptor 5 (mGluR5) activity rescues prolonged UP states in *Fmr1* KO mice ^34^. To test whether mGluR5 activity mediates the longer UP states in female Cre(+) mice, we examined the effects of a specific mGluR5 negative allosteric modulator (MTEP). Slices from Cre(+) mice or Cre(-) controls of both sexes were pretreated with vehicle or MTEP (3 µM). We confirmed a robust increase in UP state duration in female Cre(+) mice (218% of Cre(-) controls), that was reduced to normal levels by MTEP. UP state duration in Cre(-) females was unaffected by MTEP and there was a significant interaction of MTEP with genotype (F (1, 110) = 12.05; p< 0.01; Fig. 2A,B). In males, there was a small increase in UP state duration in Cre(+) mice (139% of Cre(-) controls; Fig. 2B), that was unaffected by MTEP. These results suggest a sex-specific gain of function of mGluR5 in female Cre(+) mice that drives circuit hyperexcitability.

**Figure 2.**
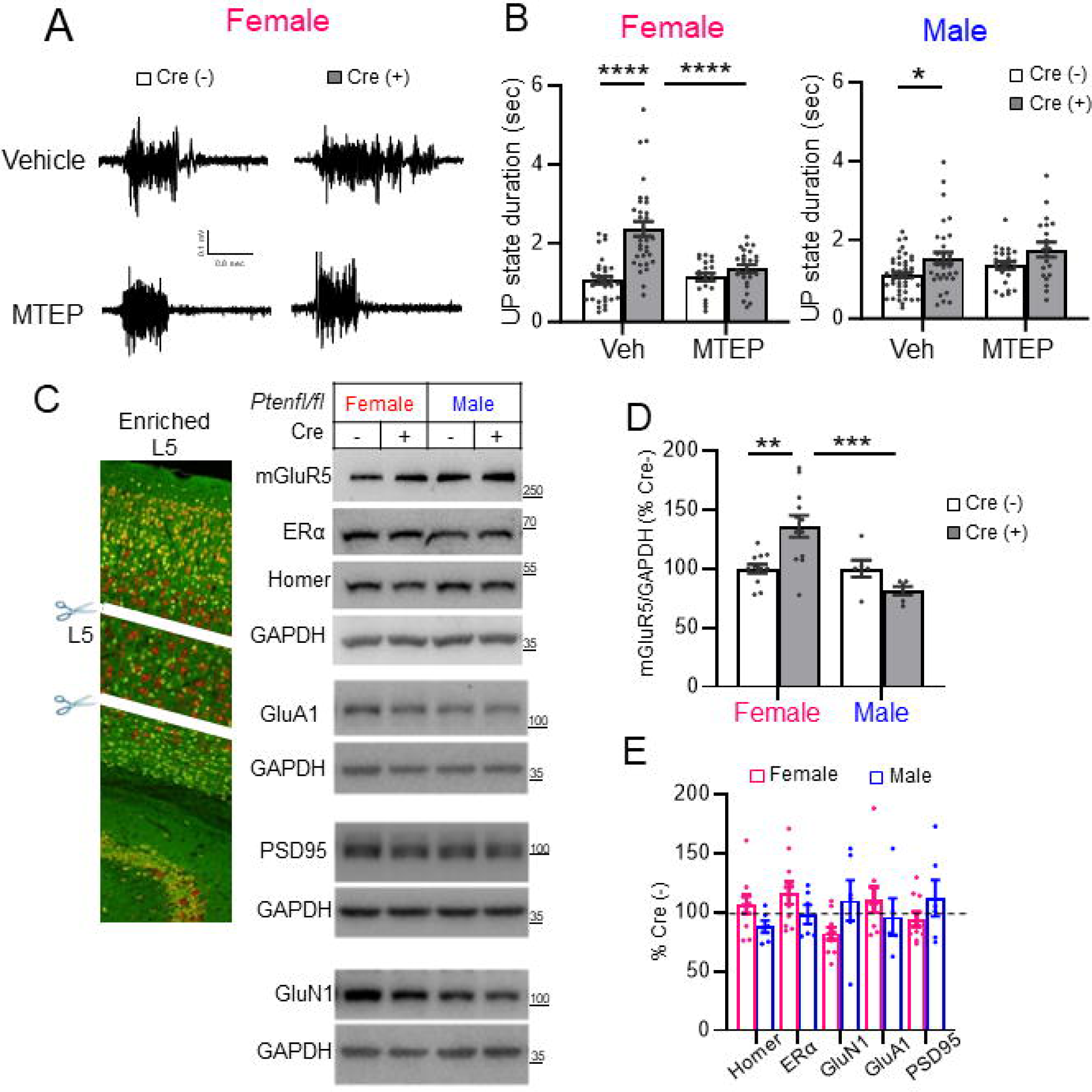
Prolonged UP states in Female ^NSE^*Pten* KO mice are rescued by an mGluR5 negative allosteric modulator (NAM) and associated with increased mGluR5 in cortical Layer 5. **A, B.** The mGluR5 selective NAM MTEP (3 μM; 2 hours) corrected UP state duration only in female Cre (+) mice without affecting UP states in Cre (-). A. Representative UP states from female Cre (-) and Cre (+) slices. **B.** Group data with N= 25-45 slices from 4-8 mice/genotype/sex. **C**, **D**. Representative western blots (C) and Quantified group data (D) show that mGluR5 levels are elevated in L5 (microdissected from slices of ^NSE^*Pten* KO mice) of female, but not male, Cre (+) mice as compared to Cre(-) same sex littermates. **E.** Quantification of mGluR5 interacting proteins, Homer and ERα, AMPA or NMDA receptors, or PSD-95 revealed no changes in L5 of Cre(+) mice of either sex.

### mGluR5 expression is selectively enhanced in female Cre(+) mice

To determine if mGluR5 levels were altered with *Pten* deletion, we performed western blotting of lysates from somatosensory cortical slices of ^NSE^*Pten* KO mice where we attempted to enrich for *Pten* KO neurons by microdissection of lower cortical layers, including L5. Western blots of cortical L5-enriched lysates revealed a small, but significant (35%) increase mGluR5 in Cre(+) female mice as compared to Cre(-) controls. mGluR5 protein levels were unaffected in male Cre(+) mice and there was a significant interaction of sex X genotype (F (1, 32) = 12.80; p< 0.01; Fig. 2C,D). The increase of mGluR5 in Cre(+) female lysates was not associated with changes in protein levels for other glutamate receptors (GluN1 or GluA1) or excitatory synaptic proteins (PSD-95 or Homer) (Fig, 2E). These data suggest a female specific upregulation of mGluR5 protein in PTEN KO neurons. In *Fmr1* KO mice, mGluR5 is less associated with its binding partner Homer, a scaffolding protein that regulates its localization to synapses, signaling to effectors and cortical circuit excitability^35^ ^36^. To determine if mGluR5 is dissociated from Homer in ^NSE^*Pten* KO mice, we performed co-immunoprecipitation of Homer and mGluR5 from lysates of either whole or L5-enriched cortical lysates from male and female mice. In contrast to *Fmr1* KO, we were not able to detect differences in mGluR5 co-IP with Homer in Cre(+) mice as compared to their same sex Cre(-) controls in either whole or L5 enriched cortical lysates (Fig. S2A,B). This suggests other mechanisms of mGluR5 dysfunction in the female ^NSE^*Pten* KO females or we are unable to detect changes with a bulk co-IP method because of the mosaic nature of Pten deletion in the ^NSE^*Pten* KO.

### Antagonism or genetic reduction of Estrogen Receptor **_α_** (ER**_α_**) in L5 neurons rescues prolonged UP states in female ^NSE^*Pten* KO mice

A large body of evidence indicates that mGluR5 functionally interacts with estrogen receptor alpha (ERα) specifically in the female brain. mGluR5 mediates the rapid non-genomic effect of ERα on multiple neuronal functions including synaptic transmission, cognition, feeding behavior and others^37–40^. Furthermore, mutations in PTEN in breast cancer cells result in enhanced ERα function through phosphorylation of ERα by either by Akt or S6K^21–23^. To determine if ERα activation contributes to circuit dysfunction in female Cre(+) somatosensory cortex we used 2 structurally distinct antagonists of ERα methyl-piperidino-pyrazole (MPP; 1µM) ^41^ ^42^ or GNE-149 ^43^. In females, MPP or GNE-149 rescued the increase in UP state duration in Cre(+) mice and had no effect on UP states in Cre(-) mice (Fig. 3A,B). There was also a significant interaction of genotype with MPP (F (1, 79) = 8.291; p< 0.01) or GNE-149 (F (1, 65) = 8.682; p= 0.01) on UP state duration. Neither MPP nor GNE-149 affected UP state duration in slices from Cre(+) male mice (Fig. S3A). These results suggest, that like mGluR5, there is a female selective signaling of ERα in Cre(+) mice that promotes UP state duration. To confirm the role of ERα in UP states using a genetic strategy and determine if increased ERα function occurred in the NSE-Cre(+) cortical neurons with *Pten* deletion, we obtained floxed *Esr1* mice and bred them to create mice with reduced ERα levels in NSE-Cre(+) and *Pten* deleted neurons (NSE-Cre/*Pten^fl/fl^/Esr1^fl/+^*) and measure UP states in these mice as compared to NSE-Cre/*Pten^fl/fl^*and Cre(-) littermates. In females, UP state duration in slices from NSE-Cre/*Pten^fl/fl^/Esr1^fl/+^*mice was reduced as compared to NSE-Cre/*Pten^fl/fl^* littermates and equal to durations measured in Cre(-) controls (Fig. 3C, D). In contrast, UP state duration in male Cre(+) mice was unaffected by genetic reduction of ERα (Fig. 3D). Heterozygosity of ERα in cortical neurons without PTEN deletion (NSE-Cre/*Pten^+/+^/Esr1^fl/+^*) had no effect on UP state duration in females (Fig. S3B). These results suggest that ERα is either overactive and/or dysfunctional in *Pten* deleted NSE-Cre(+) L5 neurons that promotes cortical circuit hyperexcitability in females, but not males. This also suggests a cell autonomous regulation of ERα by PTEN in L5 neurons that contributes to circuit dysfunction.

**Figure 3.**
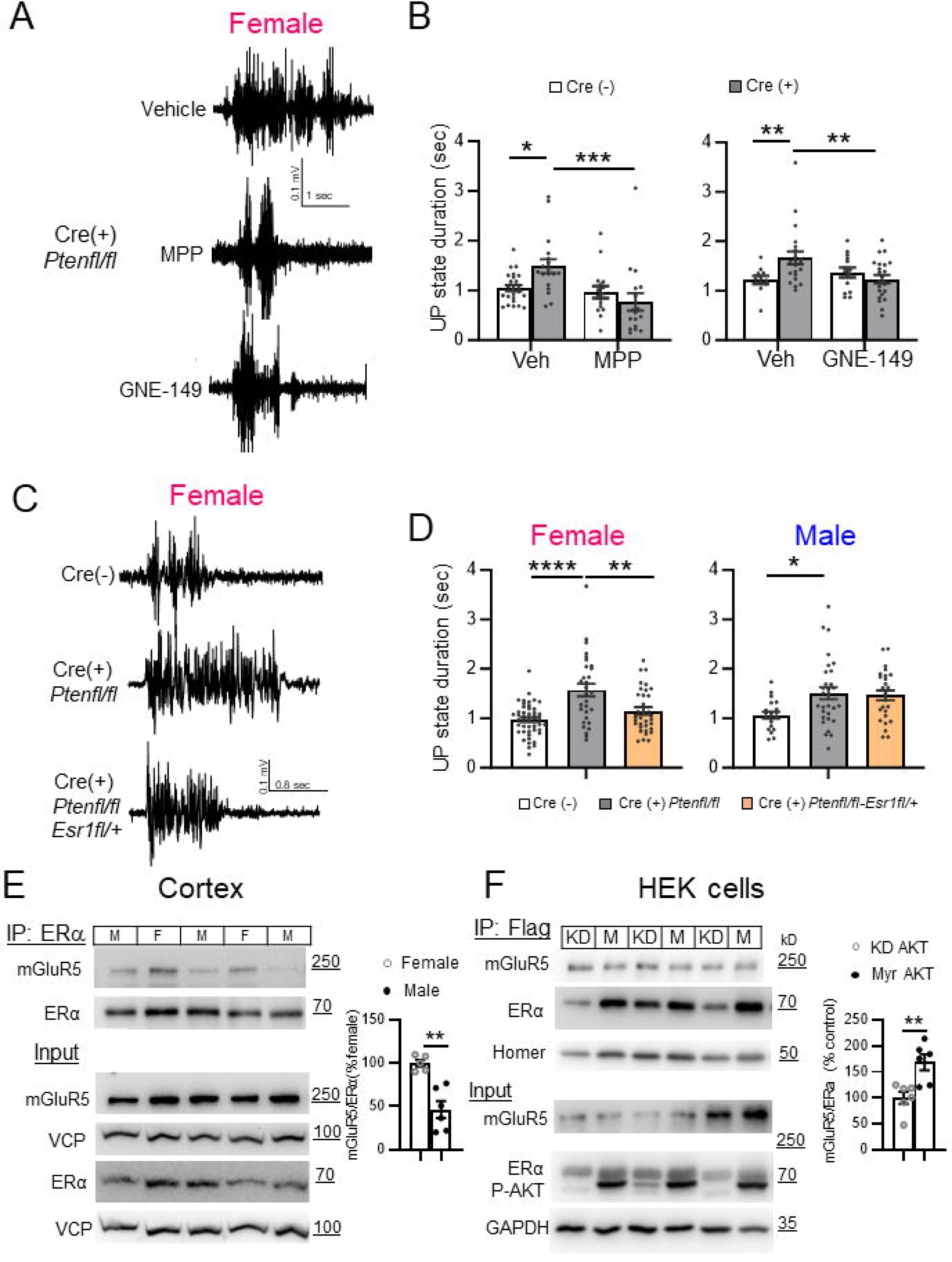
Estrogen Receptor α (ERα) is required in L5 neurons for prolonged UP states and is more associated with mGluR5 in females. **A, B.** Representative UP states (A) and group data (B) show that two ERα selective antagonist (MPP and GNE 1 μM; 1.5-2 hours) reduces UP state duration in slices from Cre (+) female mice. N=16-32 slices from 4-8 mice/genotype. **C.** Representative UP states from female *NSE-Cre/Pten^fl/fl^*(Cre (+)) and Cre(-) mice and *NSE-Cre/Pten^fl/fl^/Esr1^fl/+^*, with a genetic reduction of ERα in NSE-Cre(+) neurons. **D.** Genetic reduction of ERα expression reduces UP state duration in female (left), but not male (right), Cre (+) mice. N= 20-40 slices from 3-8 mice/sex/genotype. **E.** Left: Representative western blots (left) and quantified group data (right) show that cortical ERα-mGluR5 physical interactions (co-IP) are enhanced in females as compared to males wild-type mice (n = 5 -6). **F.** Active AKT increased association of ERα and mGluR5 in HEK cells. Representative western blots (left) and quantification (right) show co-IP of Flag-mGluR5 and HA-ERα from lysates of HEK-293T cells co-expressing either constitutively active AKT (myrAKT; M) or a kinase dead (KD) AKT. N= 6.

### mGluR5-ER**_α_** complexes are enhanced in female cortex and in response to active Akt

ERα mediates many non-genomic effects of estradiol by physically or functionally interacting with mGluR1 and mGluR5 in brain regions such as the striatum and hippocampus^39,44^. To determine if mGluR5 physically interacts with ERα in cortex and if this is regulated by sex, we performed co-IP experiments and observed an association of mGluR5 with ERα in cortical lysates that was enhanced, by ∼2-fold, in female wildtype mice as compared to males (Fig. 3E, S3C). Total mGluR5 and ERα levels were not different between sexes. In subcellular fractionation experiments, we observed that mGluR5 was present in both nuclear and non-nuclear fractions from cortical lysates and ERα was primarily non-nuclear as previously shown^45,46^(Fig. S3D). This suggests that ERα and mGluR5 interact in a non-nuclear complex in cortex. To determine if mGluR5-ERα interactions were affected by *Pten* we performed a co-IP of ERα and mGluR5 from total cortical lysates of Cre(+) and Cre(-) male and female mice (Fig. S3E,F). We observed a non-significant main effect of genotype (F (1, 20) = 4.251; p = 0.056) on mGluR5-ERα interactions, with a tendency to increase in Cre(+) mice but did not observe sex-specific effects. The mosaic nature of the NSE-Cre(+) deletion of PTEN in cortical neurons likely prohibits us from detecting molecular changes in whole cortical homogenates. We attempted to co-IP mGluR5-ERα from microdissected cortical slices to enrich for PTEN KO neurons but encountered technical challenges. To overcome challenges of working with endogenous ERα, we expressed ERα together with mGluR5 and Homer2 in HEK293 cells. As we observed in the brain, mGluR5 and ERα interact in a complex (Fig. S3G). We observed that coexpression of Homer2 increased mGluR5-ERα interaction (Fig. S3H) so we co-expressed Homer2 in this assay. To mimic the effect of *Pten* knockdown, we overexpressed a constitutively active form of AKT (myrAKT) or a kinase dead (KD-Akt; K179M)^47^. Active AKT increased association of ERα and mGluR5 (Fig. 3F). These results predict that PTEN deletion and enhanced Akt activity increases mGluR5 association with ERα.

### Acute inhibition of ERK activation or protein synthesis corrects UP state duration in female NSE-Cre PTEN KO mice

Candidate signaling pathways downstream of mGluR5 that may regulate circuit excitability include mTORC1, the mitogen-activated kinase ERK, as well as *de novo* protein synthesis ^48–55^. To test the role of these candidate pathways in prolonged UP states in female Cre(+) mice, slices were pretreated with either an inhibitor of mTORC1 (rapamycin; 200nM), or the upstream activating kinase of ERK, MEK, U0126 (20 µM) or one of two different protein synthesis inhibitors (anisomycin (20 µM) or cycloheximide (60 µM); Figs. 4A,B). Surprisingly, rapamycin pretreatment (200nM; 2 hr) had no effect on UP state duration in female Cre- or Cre+ mice and failed to correct the prolonged UP states (Fig. S4A,B). U0126 reduced and corrected UP state duration in Cre(+) females to that observed in Cre(-) controls (Fig. 4A, B). There was no effect of U0126 on UP state duration in Cre(-) females. Similarly, anisomycin and cycloheximide both corrected UP state duration in Cre(+) females and had no effect on UP states in Cre(-) controls (Fig. 4A, C, D). Thus, there was a significant interaction of inhibitor with genotype on UP state duration for both anisomycin (F (1, 57) = 5.236; p< 0.05) and cycloheximide (F (1, 55) = 8.317; p<0.01). In males, although there was no effect of genotype on UP state duration, there was a main effect of U0126 to reduce UP state duration (F (1, 51) = 6.076; p<0.05) and this was significant in Cre(+), but not Cre(-) mice (Fig. S4C). Similarly, there were main effects of anisomycin (F (1, 76) = 15.01; p<0.001) and cycloheximide (F (1, 41) = 8.443; p<0.01) to reduce UP state duration in males, but this reached statistical significance only in Cre(+), but not Cre(-), males (Fig. S4D, E). Although the UP state durations are not robustly increased in male Cre (+) mice, the UP states are sensitive to ERK and protein synthesis inhibitors. Taken together our results suggest that ERK signaling, and protein synthesis downstream signaling pathways from mGluR5 and ERα promote female selective cortical hyperexcitability in Cre(+) mice.

**Figure 4.**
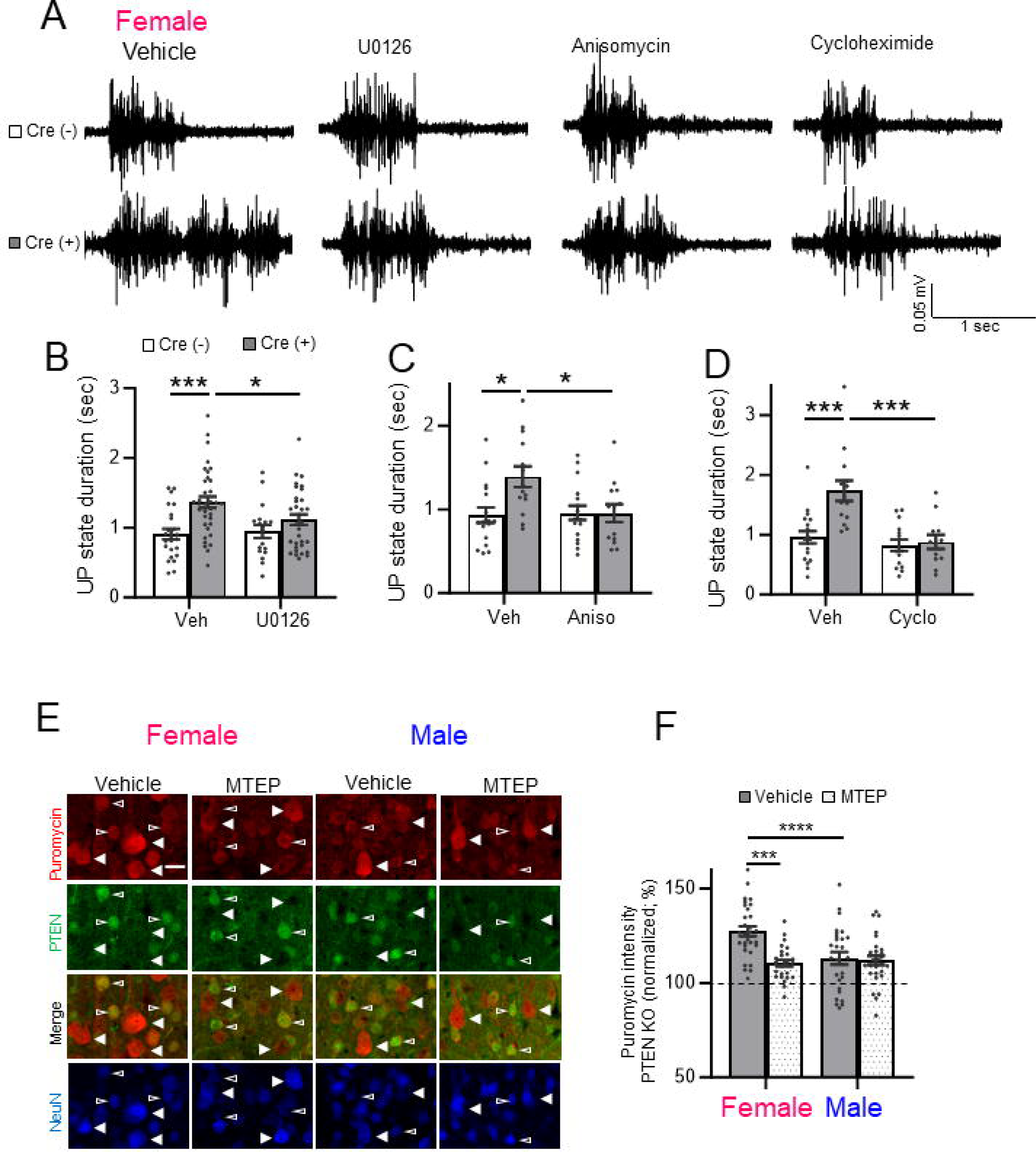
Inhibitors of ERK activation and *de novo* protein synthesis correct prolonged UP states in female ^NSE^*Pten* KO mice. **A.** Representative UP states from female Cre (-) and Cre(+) mice pre-incubated in vehicle, U126 (20 µM, 2 hrs), anisomycin (20 µM, 1hr) or cycloheximide (60 µM, 1 hr). **B, C, D**. Group averages of UP state duration in inhibitors of ERK activation or protein synthesis in female Cre (+) and Cre (-) mice. N= 12-30 slices from 3-6 mice/genotype/condition. **E.** Representative images of puromycin immunolabeling (red) of L5 neurons in cortical sections from male and female Cre (+) mice pre-treated with vehicle or MTEP (3 µM; 2 hr). NeuN (blue) and PTEN (green) immunolabeling identify PTEN KO (filled arrows) and neighboring PTEN+ neurons (open arrows). Scale bar: 30 µm. **F.** Quantified group data of puromycin fluorescent intensity of PTEN KO neurons normalized to neighboring PTEN+ neurons. N= 26-30 sections from 4 mice/sex.

From these results, we hypothesize that there is enhanced mGluR5 signaling to protein synthesis in female Cre(+) mice that promotes cortical circuit excitability. To test this hypothesis, we measured bulk protein synthesis in male and female cortical neurons with and without *Pten* deletion using surface sensing of translation (SUnSET) in the presence or absence of MTEP. SUnSET measures *de novo* translation by incorporation of the tRNA analog, puromycin^56^. Puromycin incorporation into newly synthetized protein is detected with a puromycin antibody and fluorescence immunohistochemistry (IHC) ^57^. Cortical slices from Cre(+) male and female were acutely prepared as for UP state recordings, incubated in puromycin (40 min) with or without MTEP (3µM) and processed for IHC. Neurons were identified using an antibody for NeuN or NeuroTrace (Nissl) and NSE-Cre(+), *Pten* deleted (KO) L5 neurons, were identified with a PTEN antibody. The mosaic nature of NSE-Cre expression allowed us to compare puromycin incorporation in the soma of neighboring Cre(-) or “WT” neurons, identified by positive staining for PTEN and Cre(+) or “KO” neurons that were negative for PTEN in the same section. Preincubation of slices in anisomycin strongly reduced puromycin labelling indicating that it is reflecting *de novo* protein synthesis (Fig. S4F). Analysis of puromycin fluorescence intensity revealed that PTEN KO L5 neurons in females have enhanced puromycin intensity as compared to neighboring WT neurons (127±3% of WT; n = 30 sections/4 mice) and this increase was reduced by MTEP treatment (113±3%; n = 27/4 mice; Fig. 4E, F). Using a Cre-reporter (TdTomato) to identify Cre(+) neurons, we determined that puromycin intensity is not elevated in female NSE-Cre (+) L5 neurons without *Pten* deletion (Fig. S4G, H). Therefore, the increase in protein synthesis in PTEN KO neurons is a result of *Pten* deletion and not an artifact of NSE-Cre expression. In male mice, protein synthesis, as measured by puromycin intensity, were elevated in PTEN KO L5 neurons (n=110±2% of neighboring WT, n= 26; p <0.0001; one sample t-test), but this increase was less than observed in females PTEN KO neurons (p<0.0001; Sidak’s multiple comparison) and unaffected by MTEP (n=112±2%; n=29). As a result, there was a significant interaction of MTEP and sex on puromycin intensity in PTEN KO neurons (F (1, 108) = 9.293; p< 0.01; Fig. 4F). These results indicate that mGluR5 activity drives protein synthesis in PTEN KO neurons but does so only in females. Because inhibition of ERK signaling with U0126 rescued UP state duration in female Cre(+) mice, but rapamycin did not, we tested whether U0126 or rapamycin reduced protein synthesis in PTEN KO neurons in female mice (Fig. S4I, J). Surprisingly, we observed that both U0126 and rapamycin pretreatment of slices reduced puromycin incorporation into PTEN KO neurons to a similar degree. These results suggest that inhibition of bulk protein synthesis does not correlate with rescue of the circuit hyperexcitability and suggests ERK-dependent translation of specific transcripts in PTEN KO neurons drives circuit excitability.

### Genetic reduction of ER**_α_** levels reduces protein synthesis and cell size in PTEN KO L5 neurons from female mice

Estrogen and ERα promote mRNA translation in neurons and in cancer cells ^58–60^. To determine if ERα heterozygosity reduced protein synthesis in PTEN KO neurons, we measured puromycin incorporation in neighboring WT (PTEN+) and KO (PTEN(-)) L5 neurons in the cortices of female NSE-Cre/*Pten^fl/fl^* and NSE-Cre/*Pten^fl/fl^/Esr1^fl/+^*mice (Fig. 5A, B). PTEN KO L5 neurons had a 129±3% increase in puromycin intensity as compared to neighboring WT (Cre-) neurons; n = 17 sections/3 mice. ERα heterozygosity in PTEN KO neurons reduced puromycin intensity to 118±4% of WT (n= 17; p< 0.05) suggesting that ERα contributes to the enhanced protein synthesis in female PTEN KO neurons. To determine if ERα contributed to other cellular phenotypes of PTEN KO neurons in a sex-specific manner, we measured cell size^29^. The soma size of PTEN KO L5 neurons were ∼40% larger than neighboring WT neurons in both males and females (Fig. 5C-F). ERα heterozygosity reduced the soma size of PTEN KO neurons to ∼120% of WT in females, but not had no effect on PTEN KO cell size in males (Fig. 5C-F). PTEN deletion results in activation of the mTORC1-Ribosomal S6 kinase (p70S6K) pathway, observed by phosphorylation (P) of ribosomal protein S6 (P-S6; Ser235/236; ^29^). This pathway may contribute to cell size and enhanced protein synthesis^61–63^. In female mice, we observe enhanced P-S6 in PTEN KO L5 neurons in comparison to neighboring WT neurons (Fig. S5A, B). ERα heterozygosity reduced P-S6 in PTEN KO neurons. However, the % increase in P-S6 in PTEN KO neurons as normalized to neighboring WT neurons is unchanged by ERα deletion, in contrast to protein synthesis. This may be because of the small, trending decrease in P-S6 in Cre-neurons. These results indicate that ERα contributes to the enhanced protein synthesis and cell size of PTEN KO neurons in females. Activation of P-S6 may be contributing to ERα driven protein synthesis in PTEN KO neurons, but there are likely other signaling pathways that play a role.

**Figure 5.**
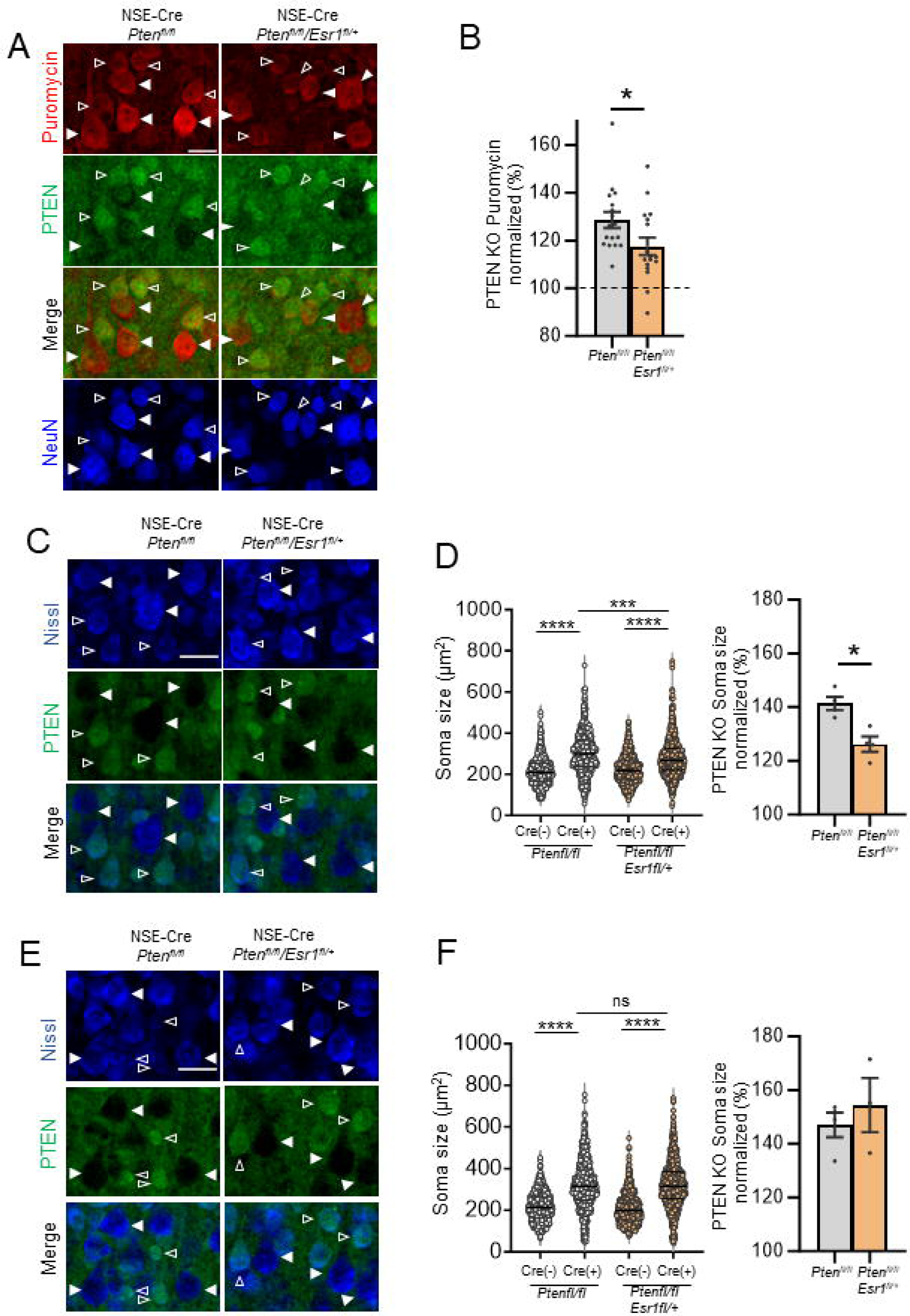
ERα promotes protein synthesis and soma size in *Pten* KO L5 neurons from female mice. **A, B.** Representative images of puromycin immunolabeling (red) of L5 neurons in cortical sections from female NSE-Cre/*Pten^fl/fl^* or NSE-Cre/*Pten^fl/fl^/Esr1^fl/+^*mice, and quantified group data of PTEN KO L5 neurons normalized to neighboring PTEN+ neurons within each genotype. N= 17 sections from 3 mice/genotype; t-test. **C, E.** Representative images of Nissl staining (blue) of L5 neurons in cortical sections from female (C) and male (E) NSE-Cre/*Pten^fl/fl^* or NSE-Cre/*Pten^fl/fl^/Esr1^fl/+^* mice. PTEN (green) immunolabeling identifies PTEN KO (filled arrows) and neighboring PTEN+ neurons (open arrows). Scale bar = 40 µm. **D, F.** Raw (left) and normalized (right) mouse averages show an increase in soma size of PTEN KO L5 neurons that is reduced by genetic reduction of *Esr1* in females (D), but not in males (F). N = 20-21 sections from 3-4 mice/sex.

Estrogen activates ERK via ERα in neurons and ERK can regulate mRNA translation through phosphorylation of eIF4E, 4E binding protein and other factors^59,64,65^. To determine if ERK signaling is selectively elevated in female PTEN KO neurons and if this relies on mGluR5 and/or ERα, we performed immunohistochemistry for phosphorylated (P) ERK (Thr202/Tyr204) in L5 neurons in sections from Cre(+) male and female mice. P-ERK levels were elevated by ∼40% in Cre(+) neurons, as compared to neighboring Cre(-) neurons in L5, in both males and females (one sample Wilcoxon test; Fig. 5C-F). Surprisingly, P-ERK levels were similarly elevated in Cre(+) neurons in slices pretreated with MTEP from either female or male mice. In contrast, antagonism of ERα with GNE-149 prevented P-ERK increases in female, but not male, Cre(+) neurons. These results suggest that ERα and mGluR5 may signal through distinct pathways in female PTEN KO neurons to promote protein synthesis and cortical circuit hyperexcitability.

### Sex specific alterations in cortical temporal processing of sensory stimuli

The prolonged cortical UP states in young female NSE-Cre PTEN KO mice suggests altered timing of cortical circuits which may impact temporal processing of sensory stimuli. High fidelity temporal processing of sound is necessary for speech and language function^66–68^ and is affected in individuals with ASD ^69–72^. Individuals with ASD show deficits in detection of sound duration, onset and offset, and rapid changes in spectrotemporal properties^73–75^. Impaired temporal gap detection thresholds in children are associated with lower phonological processing scores^70^. To determine if there are sex-specific alterations in temporal processing, particularly with rapid gaps in sounds, we performed epidural EEG recordings on the auditory (AC) and frontal cortices (FC) in young (P21-23) ^NSE^*Pten* KO mice. To address temporal acuity, we used a gap-in-noise auditory steady state response (gap-ASSR) paradigm ^76^. The gap-ASSR elicits a stead state response by presenting short gaps in continuous noise at 40 Hz. Using short gap widths and shallow amplitude modulation depths (∼75%), we compared the limits of temporal processing of cortical circuits across experimental groups. To measure the consistency of cortical responses to the gap-ASSR across trials, we calculated the inter-trial phase clustering (ITPC) at 40Hz. We also measured ITPC during the ASSR without any gaps and termed this “baseline ITPC”. Both male and female Cre+ mice showed deficits in ITPC of the gap ASSR measured in the AC and FC with a brief, 4-6ms, gap (Fig. 6A, B). However, in females the ITPC during the gap-ASSR was not greater than baseline ITPC, or when no gaps were present. Male Cre+ mice had a greater ITPC during the gap-ASSR as compared to baseline. These results suggest that female Cre+ mice are not consistently able to detect brief gaps in sound, and the cortical responses are similar to noise. Male Cre+ mice can detect the brief gaps in sound, but their responses are not as consistent as Cre-male mice. These results suggest deficits in temporal processing of sensory stimuli in both male and female Cre+ mice and these deficits are more severe in females.

**Figure 6.**
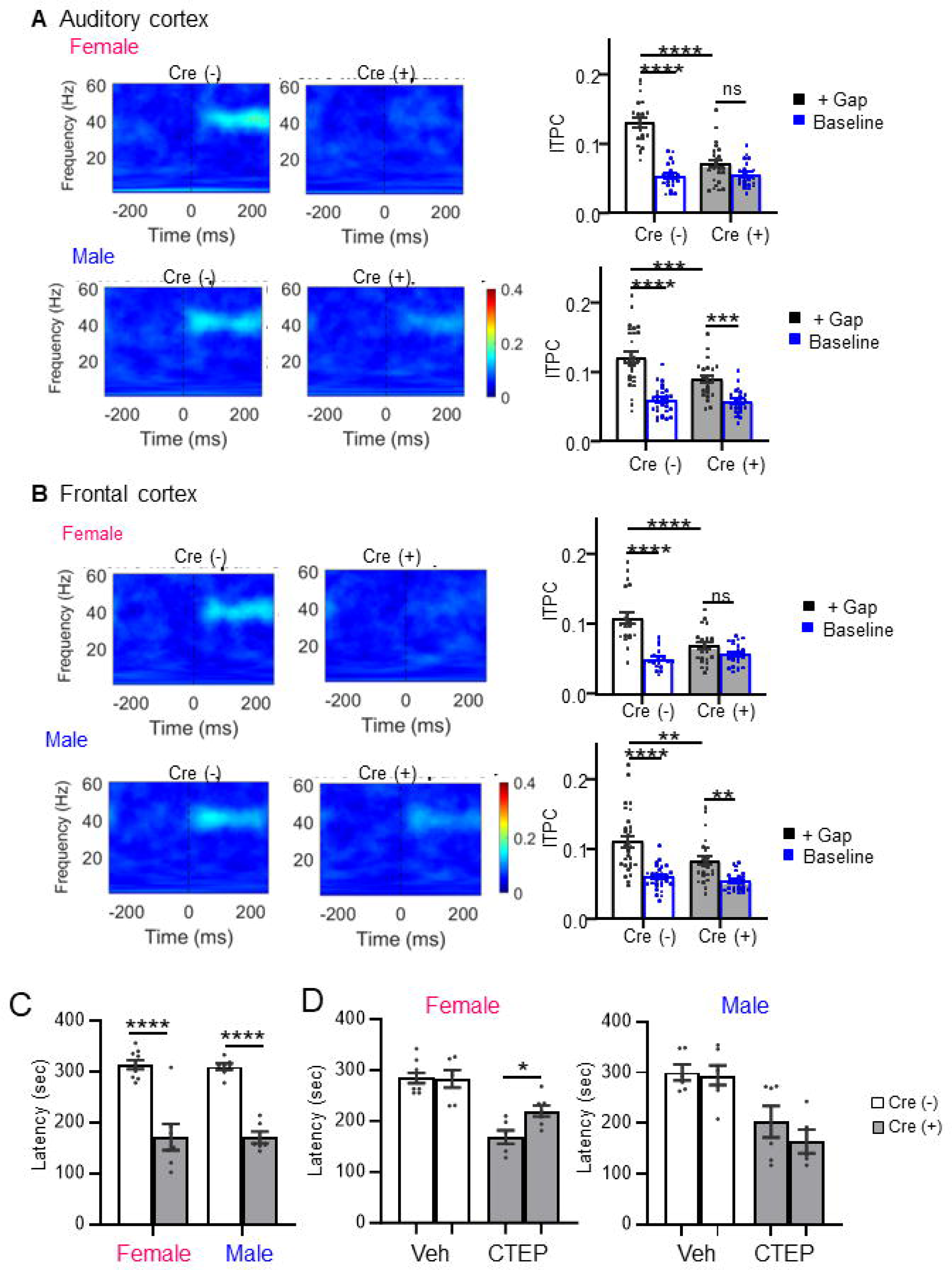
Deficits in temporal processing of sensory stimuli and mGluR5-dependent seizures female ^NSE^*Pten* KO mice. **A, B** (Left) Average ITPC heatmaps of P21 female and male Cre(-) and Cre(+) mice from the auditory cortex (A) or frontal cortex (B). Heatmaps show the average ITPC produced across presentations of the 6ms/75% modulated gap. **A, B** (Right) Both male and female Cre(+) mice show ITPC deficits when presented with gap interrupted noise (+ Gap). Cre(+) females’ ITPC during the gap-ASSR was not greater than baseline levels (unmodulated noise). Male Cre(+) mice show increased ITPC during the gap-ASSR as compared to baseline. **C.** Latency to myoclonic twitch in response to flurothyl in 6-8 weeks male and female Cre (+) and Cre (-) mice. **D.** The mGluR5 NAM CTEP increases latency to myoclonic twitch in female (left), but not male (right) Cre (+) mice. N=6-9 mice/sex/genotype/treatment.

### Seizure susceptibility in female NSE-Cre PTEN KO mice

Our slice results suggest an mGluR5-dependent hyperexcitability of cortical circuits in female NSE-Cre PTEN KO mice. To determine if there are sex-specific circuit hyperexcitability *in vivo*, we measured susceptibility to seizures induced with the volatile convulsant flurothyl. The latency to onset to distinct seizure behaviors, myoclonic twitch or tonic-clonic extension, provides a reliable index of seizure threshold in naive mice, and flurothyl seizures are highly penetrant regardless of genetic background ^77,78^. At 3 weeks of age, the age of slice for UP state experiments, neither male nor female Cre(+) mice had an increased susceptibility to flurothyl-induced seizures indicating that the cortical excitability we observe in female Cre(+) acute slices is not sufficient to affect seizure latency (data not shown). However, at 6-8 weeks of age both male and female Cre(+) mice had reduced latency to myoclonic twitch and tonic clonic seizure in response to a single flurothyl exposure (Fig. 6C). Because hyperexcitability of cortical circuits in female Cre(+) mice is dependent on mGluR5, we tested whether seizure susceptibility was also dependent on mGluR5 by administering the brain penetrant mGluR5 NAM, CTEP (i.p.; 2mg/kg), prior to flurothyl exposure. CTEP increased the latency to myoclonic twitch in female, but not in male Cre(+) mice (Fig. 6D). CTEP did not affect seizure onset latency in male or female Cre(-) mice. These results indicate that both male and female NSE-Cre PTEN KO mice develop increased susceptibility to seizures as they age, but this is dependent on mGluR5 only in females.

### Female-specific behavioral alterations in NSE-Cre PTEN KO mice

Previous work demonstrated ASD-relevant behavioral alterations in NSE-Cre PTEN KO mice such as social interaction deficits, but any sex-dependence of these deficits was not reported^29^. Female mice with a germline heterozygous deletion of *Pten* (*Pten^+/-^*) display deficits in social preference ^24,79,80^ and we hypothesized that ^NSE^*Pten* KO mice would exhibit sex-dependent behavioral deficits. To test this hypothesis, we performed a battery of behavioral tests in Cre(+) and Cre(-) male and female mice. In previous work, ^NSE^*Pten* KO mice exhibited deficits in social interaction, enhanced responses to sensory stimuli, anxiety-like behaviors, seizures, and decreased learning, which are features associated with ASD^29^. Consistent with this previous work, Cre(+) mice displayed a deficit in the three-chamber social interaction test, but this was only observed in females (Fig. S6A). Female Cre(-) mice spent more time sniffing a novel mouse in one chamber as compared to an object in another chamber, whereas female Cre(+) mice sniffed the mouse and object similarly, which resulted in a significant interaction of genotype X chamber (F (1, 42) = 7.340; p< 0.01). In contrast, male Cre(+) mice performed similarly to male Cre(-) littermates and sniffed a novel mouse more than an object (Genotype X chamber; F (1, 44) = 0.01; n.s.). Female Cre(+) mice also displayed an increase in locomotor activity, observed in dim light (females; genotype main effect; F (1, 31) = 5.874; p< 0.05; Fig. S6B). However, in a brightly lit open field, both male and female Cre(+) mice were hyperlocomotive as assessed by total distance traveled and there was no sex or genotype effect on time spent in the center of open field, a measure of anxiety (Fig. S6C). In another measure of anxiety, the dark-light box, male Cre(+) mice had decreased time spent in the light side of the box, but females also had a strong trend towards decreased time in light (Fig. S6D). These results suggest a tendency for increased anxiety in both male and female Cre(+) mice. To assess potential sex-dependence in a learning task, we assayed delay cue and context-dependent fear, or threat, conditioning. Female Cre(+) mice displayed enhanced cue-induced fear, as measured by % time freezing (Fig. S6E). Baseline freezing and context-induced freezing were normal. Male Cre(+) had normal freezing at baseline and in response to cue or context. These results suggest female specific effects of NSE-Cre mediated *Pten* deletion in mice on social preference, locomotion and fear similar to other *Pten* deletion models ^25^.

### Female specific circuit excitability in a mouse model of PTEN Hamartoma Syndrome (PTHS)

To determine if there were sex-dependent alterations in cortical excitability in a mouse model with construct validity for individuals with *PTEN* mutations, we examine UP states in acute slices of somatosensory cortex from mice with germline haploinsufficiency of *Pten* (*Pten^+/-^*). In contrast to ^NSE^*Pten* KO mice, we did not observe altered UP state duration in slices from juvenile female, or male, *Pten^+/-^* mice (P18-25; Fig. S7A). However, in young adult mice (6-8 weeks), we observed a female-specific cortical circuit hyperexcitability in *Pten^+/-^*mice (Fig. 7A, B) measured as an increase in the number of UP states and total time spent in an UP state (Fig. 7A, B). In *Pten^+/-^* males, UP state duration was reduced, and the number of UP states and total time spent in an UP state was unchanged. There was a significant interaction of genotype X sex for both UP state number (F (1, 193) = 5.041; p <0.05) and time in UP state (F (1, 193) = 7.671; p< 0.01). Surprisingly, wild-type females had fewer UP states and less time in an UP state when compared to wild-type males. These results reveal female selective increase in excitability of cortical circuits with germline haploinsufficiency of *Pten*, a human-disease relevant model of loss of function mutations in *PTEN*.

**Figure 7.**
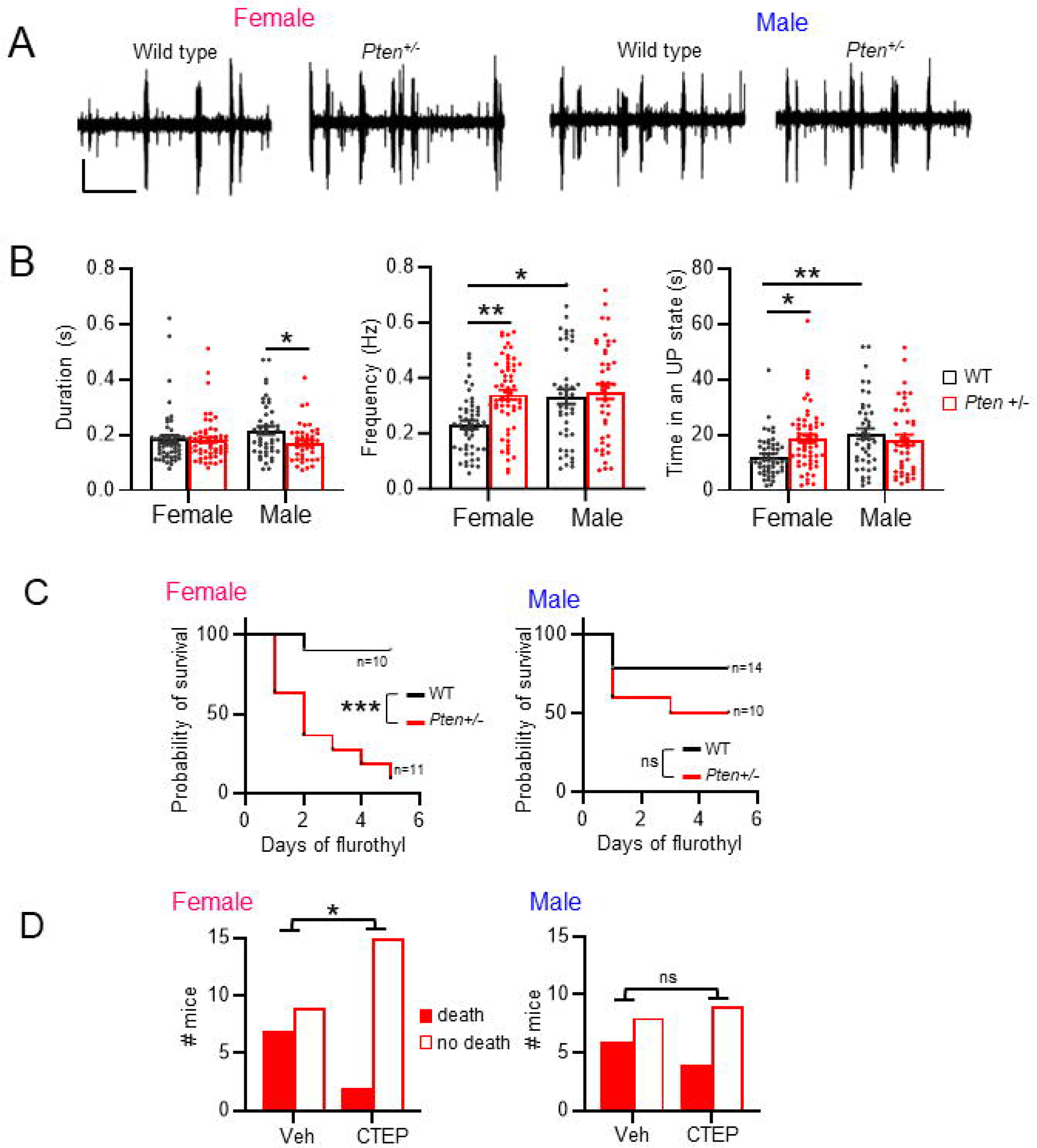
Female selective circuit excitability and mGluR5-dependent seizure severity in a mouse model of PTEN Hamartoma syndrome. **A.** Representative UP states from 6-8 week old male and female *Pten^+/-^* mice. Scale bars = 100µV/7.5sec. **B.** Group averages of UP state duration, frequency, and total time in an UP state in female and male *Pten^+/-^* slices. N= 43-58 slices from 8-12 mice/sex/genotype. **C.** Survival curve of female and male *Pten^+/-^*mice (P90-100) with repeated, daily flurothyl exposure. Mantel-Cox test. N=10-14 mice/sex/genotype. **D.** Survival of female (left) or male (right) *Pten^+/-^* mice on first day of flurothyl treatment with or without pre-treatment with CTEP. N=13-17 mice/sex/treatment condition. Chi-square test.

We next tested if there were a sex-dependent seizure susceptibility in *Pten*^+/-^ mice in response to flurothyl treatment. We observed a main effect of *Pten^+/-^* to reduce the latency for onset of myoclonic twitch and tonic clonic seizures but did not observe a female specific reduction in seizure latency (Fig. S7B). However, in a repeated flurothyl treatment paradigm (daily for 5 days), female *Pten^+/-^* mice had an increased probability of death during flurothyl treatment as compared to wild-type females (Fig. 7C). Male *Pten^+/-^* mice died at similar rate to wild-type males. To determine if the flurothyl-induced death was mediated by mGluR5, we pretreated mice daily with CTEP (i.p.) 90 min prior to flurothyl. CTEP treatment resulted in greater survival of *Pten^+/-^* females on the first day of flurothyl exposure and over the course of a 3 day repeated exposure (Fig. 7D; S7C). In contrast, CTEP did not affect survival of male *Pten^+/-^* mice. CTEP did not affect the latency to myoclonic twitch or tonic-clonic seizure in male or female *Pten^+/-^* mice and had no effect on survival of wild-type mice with repeated flurothyl (Fig. S7C, D). These results reveal a female-specific, mGluR5-dependent seizure severity in a mouse model of human *PTEN* haploinsufficiency.

## Discussion

Understanding how ASD genes affect and interact with sex-specific brain functions and signaling pathways will be key to developing more personalized therapeutics for individuals. *Pten* mutations have a strong female bias for causing breast cancer, but the sex-specific roles of PTEN in the brain are relatively unknown. Here we identify a female-specific hyperexcitability of cortical circuits and an mGluR5-dependent seizure severity in mice with postnatal deletion of *Pten* in L5 neurons or with germline heterozygous deletion of *Pten*. Using the ^NSE^*Pten* KO mouse model, we identified a female specific, cell autonomous effect of *Pten* deletion in L5 neurons that results in enhanced signaling of mGluR5-ERα to protein synthesis that drives hyperexcitability and abnormal timing of developing cortical circuits. This abnormal timing of cortical circuit correlates with reduced temporal processing of sensory stimuli in young female ^NSE^*Pten* KO mice, as measured by the gap-ASSR. Young adult ^NSE^*Pten* KO displayed female specific ASD-relevant behaviors, such as reduced social interaction, hyperactivity and enhanced fear learning. Our results reveal a molecular mechanism for the sex-specific effects of loss of function of an ASD-risk gene on neuronal and circuit function and suggest distinct pharmacological strategies, based on sex, for individuals with *PTEN* mutations and perhaps others genetic causes of ASD.

### Female selective activity of mGluR5 signaling drives excitability in PTEN KO cortical neurons

Our results suggest that either enhanced or altered signaling of mGluR5 and ERα underlies the female-specific effects of *Pten* deletion. Prolonged UP states, enhanced protein synthesis and seizure susceptibility in ^NSE^*Pten* KO mice was dependent on mGluR5 in females, but not males. Antagonism or genetic reduction of ERα corrected UP states and protein synthesis in female, but not male, ^NSE^*Pten* KO mice. Previous work in rodents has shown that estrogen signals through mGluR5 selectively in female brains and this contributes to sex-dependent effects of estrogen in drug addiction, learning and memory and other behaviors ^40,81^. Consistent with these studies, we observe enhanced mGluR5-ERα complexes in female cortex. In addition, *Pten* deletion results in a female specific enhancement of mGluR5 levels in L5-enriched cortical lysates. Together, these results point to enhanced or altered mGluR5 and ERα function or signaling in female PTEN KO neurons which drives protein synthesis. In further support of this idea, genetic reduction in ERα reduced protein synthesis and cell size in L5 PTEN KO neurons, in a cell autonomous fashion. Thus, our results suggest direct effects of PTEN on ERα function and its coupling with mGluR5 within L5 neurons. Hyperactivation of Akt in breast cancer cells results in phosphorylation and constitutive activity of ERα that further drives activation of PI3K/mTOR and tumor growth ^21–23,64^. *Pten* deletion in neurons may result in similar alterations in ERα phosphorylation and activity.

### Dysfunction of mGluR5, ERK signaling and protein synthesis: Convergence of ASD risk genes

Like female ^NSE^*Pten* KO, prolonged UP states and other measures of hyperexcitability in male *Fmr1* KO mice are dependent on mGluR5 ^34^, ERK (Fig. S4E) ^54^ and *de novo* protein synthesis ^55^ (Molinaro and Huber, unpublished). Acute antagonism of ERK activation and *de novo* protein synthesis, but, surprisingly, not mTORC1, corrected long UP states in female ^NSE^*Pten* KO slices. Although rapamycin reduced protein synthesis in PTEN KO neurons, it did not correct circuit excitability, in contrast to the inhibitor of ERK activation. Similarly, in the *Fmr1* KO, inhibition of ERK, but not mTORC1 has been implicated in circuit hyperexcitability and seizures^53,54,82^. This suggests that an ERK-dependent signaling pathway(s) promotes translation of specific mRNA transcripts that lead to circuit hyperexcitability with loss of function of either *Fmr1* or *Pten*. Although inhibitors of ERK suppressed both long UP states and protein synthesis rates in female PTEN KO neurons, P-ERK levels were similarly elevated in both male and female PTEN KO L5 neurons and unaffected by MTEP. This contrasts with the female selective increases in protein synthesis in PTEN KO L5 neurons that are mGluR5 driven. Instead, our results suggest that ERα promotes ERK signaling in female, but not male, PTEN KO neurons. Therefore, mGluR5 and ERα may stimulate distinct signaling pathways to protein synthesis in PTEN KO neurons. Genetic reduction of ERα did not correct P-S6 levels in PTEN KO neurons and may be driven by mGluR5. Additional work is needed to determine if there are altered mGluR5-ERα complexes in *Pten* deleted neurons and their signaling pathways to protein synthesis.

In *Fmr1* KO mice, mGluR5 is less associated with its scaffolding protein Homer^35,83^. Restoration of mGluR5-Homer interactions in *Fmr1* KO mice corrects prolonged UP states, seizures, and some behaviors ^36,84^. In ^NSE^*Pten* KO, we were unable to detect changes in mGluR5-Homer in either sex, which may have been because of the mosaic nature of *Pten* deletion. Alternatively, mGluR5 may be activated in PTEN KO neurons through its interaction with a hyperactive ERα. However, in both models, dysfunctional mGluR5 drives elevated mRNA translation and cortical circuit hyperexcitability. Deletion of *Tsc2,* another ASD-risk gene and suppressor of mTORC1 activation, results in mGluR5-dependent hyperexcitability or seizures in mice. The sex-dependence of this effect was either not tested^49^ or evident ^85^. As in breast cancer, one may expect dysregulation of ERα in neurons with loss of function mutations in suppressors of PI3K and sex-dependent effects on brain function and behavior.

### Developmental regulation of PTEN phenotypes and the role of estrogen

The sex-specific, ERα -dependent cortical circuit dysfunction in ^NSE^*Pten* KO was observed in young, pre-pubertal mice (P18-25), when circulating testosterone and estrogen levels are low^86^. Our results with MPP suggest that estrogen is necessary to drive ERα activity and long UP states. If so, brain synthesized estrogen may mediate ERα activation in this context. Observing sex-specific effects of *Pten* deletion young mice, before puberty, also suggests that the PTEN interacts with mechanisms of sexual differentiation of the brain; a process that occurs during the first few postnatal days in rodents ^86^. Hyperexcitability of cortical circuits was also observed in female *Pten^+/-^*but not until the young adult stage (6-8 week). This may be a consequence of the heterozygous expression of *Pten* that maintains circuit function in developing cortex. Although reductions in seizure latency were not sex dependent in ^NSE^*Pten* KO mice, seizures were sensitive to CTEP only in females. We did observe enhanced seizure severity in female *Pten^+/-^* mice, as compared to males, which we measured as an increased probability of death. This was also selectively prevented in females by CTEP. These results are consistent with a female specific, enhanced function of mGluR5 that leads to circuit hyperexcitability with *Pten* deletion. Epilepsy is more common in females with autism, as compared to males^87,88^ and brain synthesized estrogen promotes seizures ^89^. This suggests there may be roles for estrogen and/or mGluR5-ERα signaling in epilepsy in females with autism. In this study, we did not control for estrus cycle in phenotypes in adult female *Pten^+/-^* mice and would predict that estrus cycle and circulating estrogen would modulate these phenotypes.

### Sex-specific deficits in temporal sensory processing in young ^NSE^Pten KO

UP states reflect the balance of excitation and inhibition in cortical circuits and provide a measure of dynamic circuit function ^30,32^. By measuring UP states we revealed altered function of developing cortical circuits in the form of prolonged circuit activity as well as an overall increase in circuit activity in females with *Pten* deletion. The contribution of this prolonged circuit activity and hyperexcitability to *in vivo* cortical function is unknown but may contribute to the reduced ability of cortical circuits to rapidly respond and synchronize to changing sensory stimuli, as revealed here by the reduced ability of female ^NSE^*Pten* KO mice to consistently respond to brief gaps in a steady state sound. Our previous work in the Fragile X Syndrome mouse model (*Fmr1* KO) identified prolonged duration of cortical UP states. Because *Fmr1* is X-linked, we only tested males in these studies. Similar to ^NSE^*Pten* KO, cortical circuits of juvenile *Fmr1* KO mice have a reduced ability to synchronize to spectrotemporally dynamic sounds at 40-80 Hz ^90^. These results suggest similar mechanisms of cortical circuit dysfunction in developing ^NSE^*Pten* KO and *Fmr1* KO that results in prolonged circuit activity and deficits in the temporal processing of sensory stimuli. In male ^NSE^*Pten* KO, we observed smaller and less reliable increases in cortical UP state duration and smaller deficits in the gap-ASSR. These results indicate that effects of PTEN deletion are not exclusive to females but are more robust in females as compared to males. Responses to narrow gaps in sounds are commonly used measures of auditory acuity that shows prolonged maturation in children and predicts phonological aspects of speech ^70^. These data suggest that observed sex-dependent differences in gap processing may underlie abnormal speech recognition in females to a greater extent than males with *PTEN* mutations. Girls with ASD have more severe deficits in sensory processing as compared to boys, particularly in the auditory and somatosensory domains ^8^. Studies of sensory circuits and their development across sexes in ASD mouse models may contribute to understanding these deficits and developing therapies.

### Deficits in female specific behaviors in mouse models of PTEN deletion

Increasing evidence indicates that there are sex-specific effects on social behavior in multiple mouse models of PTEN loss of function ^25^. Most notably, female, but not male, *Pten^+/-^* mice have deficits in social preference^18,24,25^. Interestingly, male *Pten^m^*^3m4^ mutant mice, expressing a cytoplasmic predominant form of PTEN, exhibit enhanced sociability^26^. Consistent with these studies, we demonstrate a female-specific deficit in social interaction in ^NSE^*Pten* KO mice. *Pten^+/-^* mice have female specific alterations in fear conditioning, as we observe in the ^NSE^*Pten* KO mice ^25^. We observed a female specific hyperlocomotion under dim light, but not in the open field, where both males and females were hyperlocomotive. Modulators of mGluR5, either positive or negative, affect and correct behaviors in adult mouse ASD models caused by loss of function mutation in suppressors of mTOR or mRNA translation^91–93^ and inbred strains ^94^ but sex-specific effects were not reported. Similarly, the role of brain ERα in sex-specific ASD-relevant behaviors has not been reported in any mouse models. Modulation of mGluR5 and/or ERα may have sex-specific on behavioral phenotypes in mouse models of *Pten* deletion and other ASD models.

It is increasingly recognized that ASD manifests differently in females and males. For example, as discussed, females with autism are more likely to have epilepsy and more severe problems in sensory processing in some modalities. Here we demonstrate female selective hyperexcitability of sensory circuits, enhanced severity of seizures, and a female specific pharmacology of circuit function with loss of function of a high-impact ASD-risk gene. Importantly, we reveal roles for brain estrogen receptors in ASD-relevant phenotypes and suggests that ASD-risk genes that regulate the PI3K pathway, as in cancer, also regulate ERα in the brain and may give rise to sex-specific alterations in brain function and behavior. Our results also predict sex-specific efficacy in therapeutic strategies for individuals with *PTEN* mutations and perhaps more generally in ASD.

#### Limitations of the study

Whether the sex-specific effects of *Pten* deletion we observe in mice will be similar in human neurons or patients with *PTEN* mutations is unknown. Furthermore, we also performed most experiments with homozygous *Pten* deletion in neurons, which we think is necessary to observe robust and reproducible results in mice. Although we observed some similarities with heterozygous deletion of *Pten* in mice, whether the same effects on cortical circuit function and mGluR5 and ERα function will be observed in human neuron *Pten* heterozygous deletion or from patients is unknown.

## Resource availability

### Lead contact

Further information and requests for resources and reagents should be directed to and will be fulfilled by the lead contact, Kimberly M Huber.

### Materials availability

Plasmids generated in this study are available upon request.

## Experimental model and subject details

### Animals

Mice were group housed (5 maximum) in non-environmentally enriched cages with unrestricted food and water access and a 12 h light-dark cycle. Teklad Global 16% Protein Rodent Diet was used. This diet does not contain alfalfa or soybean meal, thus minimizing the occurrence of natural phytoestrogens.

Room temperature was maintained at 21 ± 2°C. Animal husbandry was carried out by UT Southwestern Medical Center technical staff. The animal use protocols used in this manuscript were approved by the UT Southwestern Institutional Animal Care and Use Committee and approved by the Institutional Animal Care and Use Committee at the University of California, Riverside. Congenic NSE-Cre mice ^95^ on the C57BL6 background were originally obtained from Dr. Luis F Parada (Memorial Sloan Kettering Cancer Center, New York, NY) and were crossed with floxed *Pten* (Strain #:006440). Where indicated, we also utilized the floxed *Esr1* (Strain #:032173) and Ai14 Cre-reporter mice (Strain #007914) all obtained from Jackson Laboratories (Bar Harbor, ME). Germline Pten+/− mice were generated by crossing CMV-Cre (B6.C-Tg(CMV-cre)1Cgn/J) (JAX stock number: 006054) with floxed Pten (Strain #:006440). F1 with the PTEN mutation was then bred on the C57BL/6 J background to obtain WT and Pten+/− mice.

### Cultured cell lines

The HEK293 (ATCC# CRL-1573) cell line was used for heterologous expression. Cells were grown at 37°C, 5% CO2 in 1X DMEM + Glutamax (GIBCO) containing 10% Fetal Bovine Serum and 1X Antibiotic-Antimycotic (GIBCO). Cells used in experiments were passed a maximum of twenty times. Proteins constructs were transfected or co-transfected into HEK293 using Lipofectamine 2000 reagent (ThermoFisher #11668019) based on manufacturer’s instructions.

## Method details

### Plasmids production

The N-terminal, flag-tagged mGluR5a was gifted from Dr. Robert Gereau (Washington University, St. Louis, MO) (^96^, the C-terminal HA-tagged ERα was constructed from the purchased ERα cDNA expression clone (Origene #MG227304) into BamHI and EcoRI sites of pcDNA3 vector using PCR amplification. Myc-tagged Homer2 was previously described ^97^. The myr_HA_Akt1 and myr_HA_Akt1_K179M were gifts from Dr. William Sellers (Addgene plasmids #9005 and #9006 respectively) and previously described ^47^. All constructs were verified by sequencing.

### Slice preparation and electrophysiology

Cortical slices were acutely prepared from male and female mice ages P18-60, as indicated and previously described ^34^. In brief, mice were deeply anesthetized with Ketamine/Xylazine (20mg/kg xylazine and 150mg/kg ketamine) and decapitated. The brain was transferred into ice-cold dissection buffer containing (in mM): 87 NaCl, 3 KCl, 1.25 NaH2PO4, 26 NaHCO3, 7 MgCl2, 0.5 CaCl2, 20 d-glucose, 75 sucrose, and 1.3 ascorbic acid aerating with 95% O2-5% CO2. Thalamocortical slices, 400 μm, were made on an angled block ^98^, using a vibratome Leica VT 1200S. Following cutting, slices were transferred to an interface recording chamber (Harvard Instruments) and allowed to recover for 1 h in nominal “low activity” artificial CSF (ACSF) at 32°C containing (in mM): 126 NaCl, 3 KCl, 1.25 NaH2PO4, 26 NaHCO3, 2 MgCl2, 2 CaCl2, and 25 d-glucose. Slices were then perfused with a modified “high activity” ACSF that mimics physiological ionic concentrations *in vivo*, which contained (in mM): 126 NaCl, 5 KCl, 1.25 NaH2PO4, 26 NaHCO3, 1 MgCl2, 1 CaCl2, and 25 d-glucose. Slices remained in “high activity” ACSF for 45 min prior to UP state recordings. Drugs were prepared fresh daily in vehicle and present in low and high activity ACSF or as indicated.

Spontaneously generated UP states were recorded extracellularly using 0.5 MΩ tungsten microelectrodes (FHC) placed in layer 4 of primary somatosensory cortex (S1) for most experiments or Layers 2/3 or 5, as indicated, for about 5 min/slice. Recordings were amplified 10,000-fold, sampled at 2.5 kHz, and filtered on-line between 500 Hz and 3 kHz. All measurements were analyzed off-line using custom Labview software. For visualization and analysis of UP states, traces were offset to zero, rectified, and low-pass filtered with a 0.2 Hz cutoff frequency. Using these processed traces, the threshold for detection was set at 15× the RMS noise. An event was defined as an UP state if its amplitude remained above the threshold for at least 200 ms. The end of the UP state was determined when the amplitude decreased below threshold for >600 ms. Two events occurring within 600 ms of one another were grouped into a single UP state. UP state amplitude was defined based on the filtered/rectified traces and was unitless since it was normalized to the detection threshold. This amplitude may be considered a coarse indicator of the underlying firing rates of neuronal populations. Direct measures of firing rates were not possible because individual spikes could not be isolated except during the quiet periods (the DOWN states). Data are represented by the mean ± SEM. Significant differences were determined using *t* tests, one-way ANOVA, two-way ANOVA, where appropriate (all performed with GraphPad Prism. Repeated-measures ANOVA was also used when appropriate. Sidak *post hoc* tests were performed following ANOVAs.

### Subcellular Fractionation

Postnatal day 21 mouse cortices were processed as described by mincing tissue on ice with a razor blade into 1 mm cubes. Tissue cubes or cellular pellets were resuspended in 10 volumes of Buffer “A” medium containing 2.0 mM MgCl_2_, 25 mM KCl, 10 mM HEPES (pH 7.5), and protease inhibitors (Complete Tablets; Roche Applied Science, Indianapolis, IN). After swelling for 10 min on ice, cells were homogenized in a Wheaton glass homogenizer using 15 strokes with a “B” pestle. The homogenate was filtered through 3 layers of sterile gauze and centrifuged at 1000 × *g* for 10 min. The nuclear pellet was resuspended in 3 ml of Buffer N containing 0.25 M sucrose in buffer A. Resuspended nuclei were layered over 2 ml of medium containing 1.1 M sucrose in buffer A, and then recentrifuged at 1000 × *g*. This step was repeated twice. The final nuclear pellet was resuspended in SDS sample buffer. The supernatant from the first pellet was concurrently further fractionated by centrifugation at 35,000 × *g* for 40 min. This second supernatant represented soluble, cytoplasmic proteins whereas the high-speed pellet contained plasma membrane proteins. Aliquots from each fraction were used for gel electrophoresis as well as membrane binding. Protein concentrations were determined using the BCA protein assay kit (Pierce).

### Western blotting

Western blotting was performed on lysates from somatosensory cortex, or microdissected lower cortical layers (to enrich for *Pten* KO neurons), *subcellular fractions from* P21 mouse cortices, and eluted immunoprecipitated proteins. With the exclusion of the immunoprecipitated protein, sample were lysed with 50 mm Tris, pH 7.4, 120 mm NaCl, 1 mM EDTA and 1% Triton X-100, containing Protease Inhibitor Mixture, (Sigma P8340); and phosphatase inhibitor mixture 2 and 3, (Sigma P5726 and P0040). Samples were further homogenized using brief (5–10 s) pulses of sonication until lysates were clear. Lysates were then centrifuged at 10,000 g for 10 min at 4°C, to remove insoluble material. Protein concentration was determined by a BCA protein assay kit (Pierce). To equalize protein concentrations, samples were diluted with lysis buffer and 4x SDS-Sample buffer to a final concentration of 1 to 3 µg/µl. Equal amounts of protein (5–15 μg) in an equal volume were loaded for each sample on 8-10% Tris-glycine polyacrylamide gels; the gels were run using Tris-glycine SDS running buffer at constant voltage and then transferred to a PVDF membrane. Membranes were cut at appropriate molecular weights to allow immunoblotting of the same membrane for multiple proteins. After blocking with 5% milk in 1× TBS, 0.05% Tween 20 for 1 h, membranes were incubated with primary antibodies in blocking buffer overnight at 4°C. Following incubation in primary antibody overnight at 4°C, immunoblots were incubated with HRP-conjugated secondary antibodies in 5% milk in 1× TBS, 0.05% Tween 20 for 1 h at room temperature. Membranes were washed and developed using the SuperSignal™ West Pico PLUS Chemiluminescent Substrate (34580, ThermoFisher). Signal was acquired using Chemidoc MP (BioRad) and analyzed using ImageLab (BioRad). Protein bands were normalized using GAPDH,VCP or actin as housekeeping proteins. The details of the primary antibodies used, and species they were raised in, are provided in the key resources table.

### mGluR5 Co-Immunoprecipitation in vivo

Neocortices from P21 male and female ^NSE^*Pten* KO and WT mice were lysed in coimmunoprecipitation (Co-IP) buffer (50 mM Tris, pH 7.4, 120 mM NaCl, 1 mM EDTA, 1% Triton X-100 containing Protease and Phosphatase inhibitor mixture, and further homogenized using brief (5–10 s) pulses of sonication until lysates were clear. After homogenization, lysates were rotated for 30 minutes at 4°C followed by centrifugation at 10,000 X g for 10 minutes at 4°C. Then 10% of the lysate was set aside for use as an input control and the rest was tumbled overnight at 4°C with 7 µl of ERα antibody or 1.4 µg of Homer antibody (Santa Cruz Biotechnology, D-3). The same isotype IgG was used as control for the specificity of the ERα (Abcam #ab172730) and Homer D3 antibodies. 25 µl of Dynabeads Protein A or G slurry (Invitrogen) were washed in co-immunoprecipitation buffer and added to the samples for 3 h at 4 °C with gentle rotation and the beads were then washed with Co-IP buffer. Immunoprecipitated proteins were eluted with 2x SDS Sample Buffer (Bio-Rad). Eluted immunoprecipitated proteins were run in parallel with input lysate. Western blotting was performed with antibodies against Homer 1 (Santa Cruz, E-18), mGluR5, and ERα. The TrueBlot secondary Ab (18-8816-31Rockland) was used as secondary for ERα detection. For analysis of each experiment, the Co-IP band intensity values were normalized by the intensity of the corresponding IP protein band. (e.g. mGluR5/ERα or mGluR5/Homer).

### Co-Immunoprecipitations in HEK Cells

Proteins constructs were transfected or co-transfected into HEK293 using Lipofectamine 2000 reagent. 48 hours post-transfection, cells were lysed in Co-IP buffer (50 mm Tris, pH 7.4, 120 mm NaCl, 1 mM EDTA, 1% Triton X-100 containing Protease Inhibitor Mixture, and phosphatase inhibitor mixture 2 and 3, and insoluble debris was removed by centrifugation (10 min, 10,000 g). The cleared lysate was incubated with an anti-Flag antibody (Sigma-Aldrich, F1804) or same IgG isotype control, while rotating, overnight at 4°C and then for 3-4 hours with Dynabeads G magnetic beads. Bound proteins were eluted with sample buffer and loaded on gradient SDS-PAGE gels and processed for western blot as described above. Input lysate and Co-IP complexes were blotted with anti-mGluR5 and anti-ERα antibodies.

### Surface Sensing of Translation (SUnSET) in Brain Slice

To measure protein synthesis in L5 of S1 somatosensory cortex we used the immunohistochemical version of SUnSET ^56^. Acute slices were prepared as for UP states and recovered for 2 hours (1 hour in low activity ACSF and then 1 hour in high activity ACSF) in a submerged chamber with continuous aeration with 95% O2/ 5% CO2. Puromycin (2.0 µg/µl) (P8833, Sigma-Aldrich Inc.) was then added to ACSF for 40 min. MTEP (3 µM), rapamycin (200 nM) or U0126 (20 µM) were present during the 2 hours recovery time and during the puromycin incubation. For anisomycin (20 µM) control experiment, slices were incubated with the protein synthesis inhibitor for 60 min before puromycin incubation. At the end of puromycin incubation, slices were washed in cold ACSF and then fixed for 3 hours in 4% PFA, rinsed in PBS and stored at 4°C. Slices were then re-sectioned (50 µm thickness) using a Leika 1000S vibratome and stored in anti-freezing solution (30 % Glycerol, 30% Ethylene Glycol, 40 % PBS) at -20°C until subjected to immunohistochemistry.

### Immunohistochemistry

For PTEN, Phospho-S6 immunohistochemistry and soma size measurement (Fig. 5), mice were anesthetized with Ketamine/xylazine and transcardially perfused with phosphate-buffered saline (PBS) followed by 4% paraformaldehyde (PFA) in 0.01 M phosphate buffer. Brains were then post-fixed overnight in 4% PFA. After postfix, brains were washed with PBS and sectioned into 50 µm thick coronal sections using a vibratome (Leica VT 1000 S). For P-ERK and Puromycin staining after SUnSET, post-fixed slices were washed in 0.01 M PBS and re-sectioned into 50 µm thick sections containing the somatosensory cortex using a vibratome. Sections containing the somatosensory cortex were permeabilized with 0.3% Triton X-100 for 30 min, blocked with 0.1% Triton X-100, 5% goat serum, and then incubated for 24 to 48 hours at 4°C with the following primary antibodies: rabbit anti-PTEN, rabbit anti-Phospho-S6 Ribosomal Protein (Ser235/236) Alexa Fluor 647, anti-puromycin, anti-Phospho-p44/42 MAPK (Erk1/2) (Thr202/Tyr204) and using an anti-NeuN or using NeuroTrace™ 435/455 Blue Fluorescent Nissl Stain (Invitrogen). To label neurons and for soma size analysis. After the incubation in primary antibodies, sections were washed in PBS 0.04% Triton X100 at room temperature (3 × 10 min). Following the last wash, slices were incubated in secondary antibody for 2 h at room temperature, followed by 3 × 10 min washes in PBS-Triton at room temperature. Secondary antibodies used were the following: goat Anti-Rabbit IgG fluorescein (FITC)), goat anti-Mouse Texas Red-X (Invitrogen, Catalog # T-6390), Goat anti-Guinea Pig IgG (H+L) Highly Cross-Adsorbed Secondary Antibody, Alexa Fluor™ 647 (Invitrogen, Cat. N.: Cat # A-21450). NeuroTrace™ 435/455 Blue Fluorescent Nissl Stain was incubated with secondary antibodies. For P-S6 staining, PTEN stained sections (primary + secondary and Neurotrace) were incubated with the anti P-S6 antibody Alexa Fluor^®^ 647 Conjugate overnight at 4°C. Following wash with PBS 0.04% TritonX-100, sections were mounted using Aqua-Poly/Mount (Polysciences) on glass slides and imaged. ERK phosphorylation was evaluated with immunohistochemistry. Slices were permeabilized with 0.3% Triton X-100 and blocked with 0.1% Triton X-100, 5% goat serum. The Streptavidin /Biotin blocking Kit (Vector Laboratories) was also used as directed by the manufacturer. Sections were incubated with a primary antibody for Phospho-p44/42 MAPK (Erk1/2) (Thr202/Tyr204) (Cell Signaling) for 48 hours at 4°C. After that, the sections were washed and incubated first with a Biotinylated Goat Anti-Rabbit IgG Antibody (over-night at 4°C-Vector Laboratories), and then with Alexa Fluor™ 594 streptavidin (3 hours – room temperature-Thermo Fisher). After extensive washing, slices were incubated again with a rabbit antibody raised against PTEN over-night at 4°C (Cell Signaling). After washing, the sections were then incubated with a Goat antirabbit IgG conjugated with Alexa Fluor-647 (Thermo Fisher Scientific) and NeuroTrace™ 435/455 blue fluorescent Nissl stain (Thermo Fisher Scientific) to identify neuron’s soma (Thermo Fisher Scientific). This combination of reagents prevents nonspecific binding and overlap between the P-ERK and the PTEN staining (data not shown).

### Confocal Microscopy and Image quantification

Fluorescence images (1024 × 1024 pixels, 8 bit) were taken from L5 of S1 cortex using a Zeiss LSM 710 laser-scanning confocal microscope with a 20× objective, 1× zoom, 2 μm z-step and a resolution of 2.41 pixels/µm. For all quantitative comparison experiments, the same microscope and acquisition settings were used for each image and samples were processed in batches to include matched control and experimental samples. All images were processed using Fiji software(https://imagej.net/software/fiji/). For image analysis, the investigators were blind to the treatment condition.

To quantify puromycin, P-S6 and P-ERK fluorescence intensity and soma size, background was subtracted using the “Subtract background” function on FIJI (rolling ball radius: 50 pixel). After background was subtracted from each image, regions of interest (ROIs) were manually drawn around neuronal bodies in the puromycin Channel (red Channel), P-S6 channels (ultrared channel) and Neurotrace channel (blue Channel) of each image with the “Freehand” tool and were then used to measure area and fluorescence mean gray value in the Puromycin, P-S6, P-ERK, Neurotrace or NeuN channels. Non neuronal cells were excluded from the analysis based on the NeuN (for the Puromycin IHC) or Nissl staining. Cre (+) or Cre (-) neurons were then identified and separated based on PTEN immunofluorescence (Green Channel). The fluorescence intensity and area of 30 to 60 Cre (-) and (+) neurons was measured in each section for Puromycin, P-S6, P-ERK and soma size evaluations. The fluorescence intensity and area of all Cre (-) and Cre (+) neurons was averaged for each section and the mean value of the fluorescence intensity of Cre (+) was expressed as % of the Cre (-) neurons for each channel. 5-8 sections were analyzed from 4 mice for each condition/treatment. N for these data represent the # of sections. To quantify PTEN (+) and PTEN (-) neurons in Fig. S1, NeuN+ neurons were counted in FIJI by an author blinded to the sex within a L5 cortical region and then cells subsequently defined as PTEN-positive or negative.

### Flurothyl induced seizures

Flurothyl (bis(2,2,2-trifluoroethyl) ether) seizure experiments were performed as described ^100^. Each mouse was placed in a 2 liter glass chamber inside of a chemical fume hood and allowed to habituate for 1 min before the top of the chamber was closed. 10% flurothyl (bis-2,2,2-trifluoroethyl ether; Sigma-Aldrich) in 95% ethanol was then infused at a rate of 200 µL/min onto a disk of filter paper (Whatman, Grade 1) suspended at the top of the chamber. Mice exhibit various stages of increasing seizure severity in response to flurothyl exposure, including myoclonic seizure (sudden involuntary jerk/shock-like movements involving the face, trunk, and/or limbs) and generalized seizure (also known as clonic-forebrain seizures that are characterized by clonus of the face and limbs, loss of postural control, rearing, and falling). Generalized seizures can immediately progress into brain stem seizures manifested by tonic extension of the limbs. Upon emergence of a generalized seizure, the lid of the chamber was immediately removed, allowing for rapid dissipation of the flurothyl vapors and exposure of the mouse to fresh air. Mice were then returned to their home cage following recovery from behavioral seizures. One mouse at a time was tested in the flurothyl chamber, which was recharged with fresh filter paper, cleaned using water, and thoroughly dried between subjects. In the same experiments, flurothyl exposures were repeated once daily over five consecutive days. Mouse behavior during each flurothyl exposure was video-recorded and viewed by investigators (blind to genotype) who determined latency to the onset of myoclonic and generalized seizures.

The mGluR5 specific negative allosteric modulator CTEP (chloro-4-((2,5-dimethyl-1-(4-(trifluoromethoxy)phenyl)-1H-imidazol-4-yl)ethynyl)pyridine) was prepared fresh daily as a microsuspension in vehicle (0.9% NaCl, 0.3% Tween-80) and administered by intraperitoneal (i.p.) injection 90 min prior to fluorothyl exposure.

#### Behavioral tests

Behavior tests were performed during the light phase of light/dark cycle. The battery of tests was performed in the same order by all mice (-Locomotor Activity-Open Field-Dark Light Box-Three-chamber social approach-fear conditioning) and designed to minimize carryover effects ^101^. Mice were moved into the testing area at least 1 h prior to testing. Manual scoring for social recognition and fear conditioning (with a stopwatch)) was performed by a trained observer blind to sex and genotype.

### Locomotor Activity

To assess locomotor activity, mice were placed in a new home cage with minimal bedding for 2 hr and horizontal activity was monitored using photobeams linked to computer data acquisition software (San Diego Instruments) under red light condition.

### Open field test

The open field test was used to evaluate the voluntary behavior, exploratory behavior, and anxiety of experimental animals in new and different environment^102^. The light intensity in the open field apparatus was 60 lux. At the beginning of the test, the mice were placed in a PVC box (45LJ×LJ45LJ×LJ30 cm) at the same starting position and an overhead camera and tracking software (Noldus Ethovision software version 12.5) was used to monitor the mouse movement for 10 min. Total distance and time traveled in different compartment of the chamber (center, non-periphery, periphery) were measured. The chamber was cleaned after each trial. All behavioral results were analyzed by investigators blind to genotypes.

#### Light–dark (L/D) test

The apparatus comprised of a box divided into two separate compartments. The light chamber was painted white, while the other chamber was entirely black and enclosed. The L/D chambers were separated by a sliding door. After 2 min of habituation in the dark chamber the door to the light chamber was opened and mice were allowed to explore for 10 min. Behavioral parameters such latency time to leave the dark chamber and total time spent in the light and dark chamber were recorded^103^.

#### Three chamber sociability test

Social preference of a mouse was evaluated in a three-chamber choice task^104,105^. The sociability test was used to evaluate a mouse’s preference for a social object over a non-social object. All tests were conducted in a white opaque acrylic three-chambered apparatus. The apparatus was divided into three chambers connected by doorways under white light conditions. During the habituation phase, the test subject was acclimated for 10 min in the center chamber, followed by another 10 min exploring the entire apparatus. Then, the social interaction test was conducted. The test subject was first confined to the center chamber with both doorways closed. An empty wire cage was then placed in one side chamber and an identical wire cage containing an unfamiliar, sex-matched mouse (Stranger #1) was placed in the other side chamber. The doorways were then re-opened, and the test subject was allowed to freely explore the testing chambers for 10 min. Time spent directly interacting with the novel animal and novel object (sniffing time) was scored with a stopwatch by an examiner who was blind to genotype. Animals were tested between 10 and 14 weeks of age. Light at the center of the three-chambered apparatus was 60 lux for all experiments.

#### Fear Conditioning

Fear conditioning was measured in boxes equipped with a metal grid floor connected to a scrambled shock generator (Med Associates Inc., St. Albans) as described^105^. For training, mice were individually placed in the chamber. After 2 min, the mice received 3 tone-shock pairings (30 sec white noise, 80 dB, co-terminated with a 2 sec, 0.5 mA footshock, 1 min intertrial interval). The following day, memory of the context was measured by placing the mice into the same chambers and freezing was measured automatically by the Med Associates software for 5 min. Forty-eight hours after training, memory for the white noise cue was measured by placing the mice in a box with altered floors and walls, different lighting, and a vanilla smell. Freezing was measured for 3 min, then the noise cue was turned on for an additional 3 min and freezing was measured.

### EEG recordings

Mice underwent epidural electrode implant surgery at P18-20. Mice were anesthetized using i.p. injections of 80/20 mg/kg of ketamine/xylazine and were monitored closely throughout the procedure by toe pinch reflex every 10-15 minutes. ETHIQA-XR (1-shot buprenorphine, 3.25 mg/kg body weight) was administered (s.c.) prior to surgery to ensure an adequate analgesic level was present post-surgically. Following the removal of fur and skin, and sterilization (alcohol and iodine wipes) of the scalp, an incision was made to expose the scalp. A Foredom dental drill was utilized to drill ∼1mm diameter holes in the skull over the right auditory cortex (AC), right frontal cortex (FC), and left occipital cortex. The screw positions were determined using skull landmarks and coordinates previously reported ^106^ and were based on single unit mapping ^107^. The wires extending from three-channel posts were wrapped around 1 mm screws and screwed into holes. Dental cement was applied around the screws, on the base of the post, and exposed skull, to secure the implant. Mice were placed on a heating pad until fully awake and were allowed 48-72 hours for recovery before EEG recordings were made. Mice were housed individually following surgery. All EEG recordings were obtained from awake and freely moving mice. EEG recordings were performed at P20-23. Recordings were obtained from the AC and FC electrodes, using the occipital screw as reference. Mice were placed in an arena where they could move freely during the recording. The arena was inside a Faraday cage placed on a vibration isolation table in a sound-insulated and anechoic booth (Gretch-Ken, OR). Mice were briefly anesthetized with isoflurane and attached to an EEG cable. The EEG recording set-up has been previously reported ^76^. Briefly, the attached cable was connected via a commutator to a TDT (Tucker Davis Technologies, FL) RA4LI/RA4PA headstage/pre-amp, which was connected to a TDT RZ6 multi-I/O processor. OpenEx (TDT) was used to simultaneously record EEG signals and operate the LED light used to synchronize the video and waveform data. TTL pulses marked stimulus onsets on a separate channel in the collected EEG data. The EEG signals were recorded at a sampling rate of 24 kHz and down-sampled to 1024 Hz for analysis. All raw EEG recordings were visually examined prior to analysis for artifacts, including loss of signal or signs of clipping, but none were seen.

#### Gap-ASSR

The stimulus used to assess auditory temporal processing is termed the ‘40 Hz gaps in noise-ASSR’ (auditory steady state response, henceforth, ‘gap-ASSR’) ^76^ and contains alternating 250 ms segments of noise and gap interrupted noise presented at 75 dB SPL. The gaps are placed 25 ms apart, resulting in a repetition rate of 40 Hz. The 40 Hz modulation rate produces the strongest ASSR signal across temporal and frontal cortex and may reflect the resonance frequency of the underlying neural circuits ^108,109^. The gap width and modulation depth are chosen at random for each gap-in-noise segment. For data shown in Fig. 6, gaps included 4 or 6 ms and modulation depths of 75%. To measure the ability of the cortex to consistently respond to the gaps in noise, inter-trial phase clustering (ITPC) at 40 Hz was measured ^110^. The EEG trace was transformed using a dynamic complex Morlet wavelet transform. The trials corresponding to each parametric pair (gap duration + modulation depth) were grouped together. The ITPC was calculated for each time-frequency point as the average vector for each of the phase unit vectors recorded across trials (trial count >100 trials per parametric pair). The ITPC values at 40Hz were averaged to extract the mean ITPC for the parametric pairs in the AC and FC. To ensure that the calculated ITPC was not due to spurious or transient phase locking, we measured ‘baseline’ ITPC. Baseline ITPC was calculated during the periods of unmodulated noise (no gaps) and compared to the ITPC generated during periods of modulated noise.

#### Statistical Analysis

Whenever possible, data was analyzed by an investigator blind to the genotype or sex of mice. Results in figures are graphically represented as mean ± SEM. Statistics were performed in GraphPad Prism. Normal distribution was tested by the D’Agostino & Pearson test. For 2-group comparison, we used a student unpaired 2-tailed t test, or a Mann-Whitney nonparametric test. For multiple group comparison, data were analyzed using ordinary 1-way ANOVA or Kruskal-Wallis (nonparametric) or 2-way ANOVA where appropriate. Sidak and Bonferroni posthoc multiple comparison tests were used.

## Supporting information

Supplemental informations

## Credit authorship contribution statement

Gemma Molinaro: Conceptualization, Investigation, Formal analysis, Writing – original draft. Jacob E. Bowles: Investigation. Katilynne Croom: Investigation, Formal analysis. Darya Gonzalez: Investigation. Saba Mirjafary Investigation. Shari Birnbaum: Conceptualization, Supervision. Khaleel A. Razak: Conceptualization, Supervision, Formal analysis. Jay R. Gibson: Conceptualization, Supervision, Validation. Kimberly M. Huber: Conceptualization, Formal analysis, Writing – original draft, Supervision, Funding acquisition.

## Acknowledgements

This work was supported by NIH grants R37NS114516 to K.M.H. We would like to thank Dr. Luis Parada for providing the NSE-Cre mouse line and Courtney Saqueton and Christopher Williams for technical assistance.

