## Supplemental informations for "Female specific dysfunction of sensory neocortical circuits in a mouse model of autism mediated by mGluR5 and Estrogen Receptor α"

Supplementary Table 1. UP state amplitude and frequency are not different between genotype or sex.

| | | Female | | Male | |
|---|---|---|---|---|---|
| | | Cre - | Cre+ | Cre - | Cre+ |
| Layer 4 | | | | | |
| | Duration (ms) | 1014±109 | 1585±116** | 1223±127 | 1355±158 |
| | Amplitude (mV) | 0.13±0.01 | 0.14±0.01 | 0.16±0.01 | 0.13±0.01 |
| | Frequency (Hz) | 0.13±0.01 | 0.15±0.01 | 0.09±0.01 | 0.10±0.01 |
| | n (# slices) | 14 | 22 | 18 | 14 |
| Layer 2/3 | | | | | |
| | Duration (ms) | 934±83 | 1790±181*** | 1157±104 | 1301±192 |
| | Amplitude (mV) | 0.17±0.02 | 0.17±0.02 | 0.19±0.02 | 0.18±0.01 |
| | Frequency (Hz) | 0.15±0.01 | 0.13±0.01 | 0.13±0.01 | 0.15±0.01 |
| | n (# slices) | 17 | 19 | 21 | 20 |
| Layer 5 | | | | | |
| | Duration (ms) | 1032±81 | 1969±208** | 1425±153 | 1437±179 |
| | Amplitude (mV) | 0.18±0.02 | 0.25±0.03 | 0.12±0.03 | 0.19±0.02 |
| | Frequency (Hz) | 0.17±0.02 | 0.14±0.01 | 0.13±0.01 | 0.14±0.01 |
| | n (# slices) | 17 | 20 | 17 | 21 |
| 2 way ANOVA; Sidak's posthoc multiple comparison Cre- vs Cre+ within sex; **p<0.01; ***p< 0.001; | | | | | |

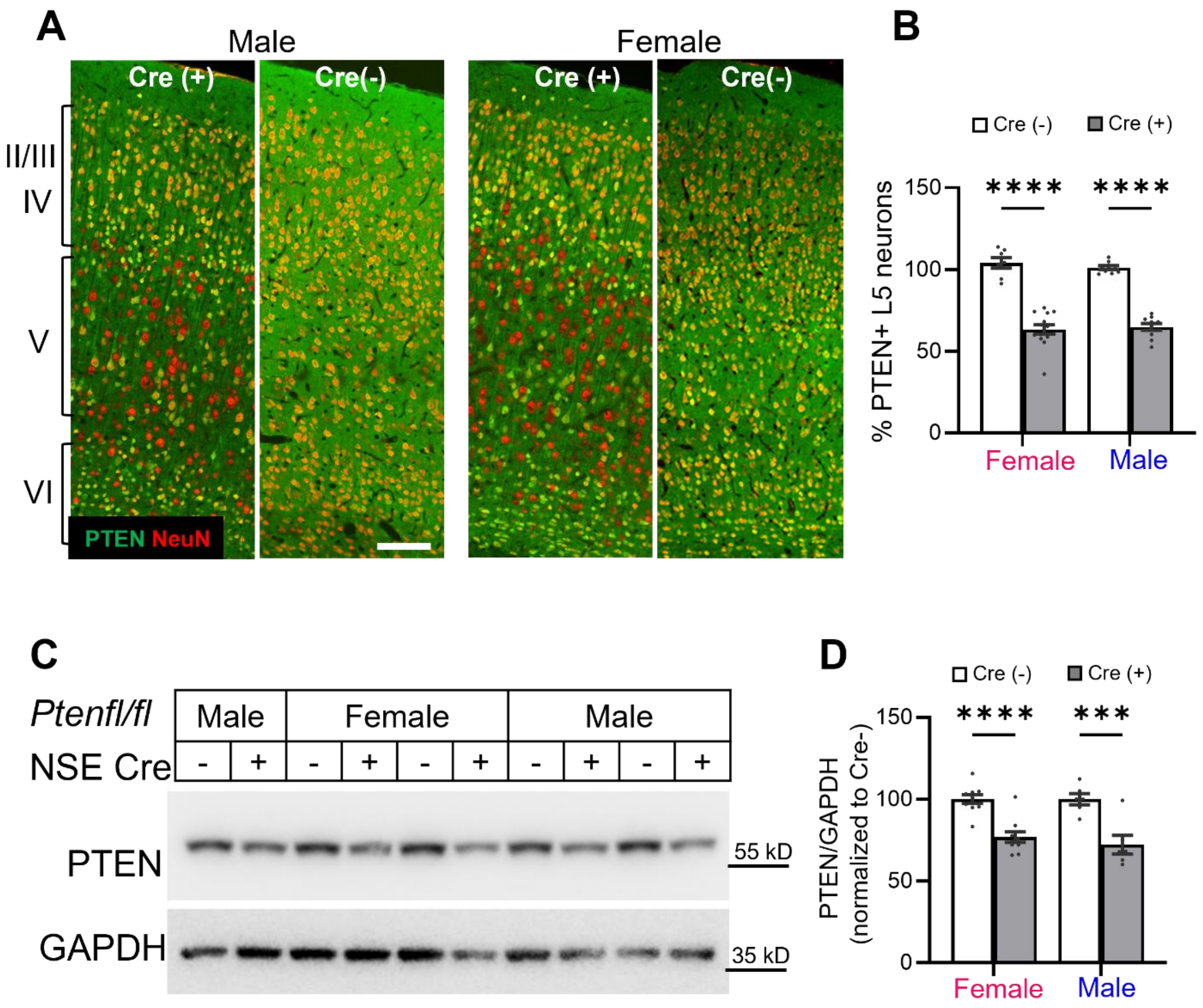


**Supplemental Figure 1. The female specific effects on cortical circuit function are not attributable to sex-dependent differences in PTEN levels in $^{NSE}$*Pten* KO mice. A.** Representative images of immunohistochemistry for PTEN (green) and the neuronal marker NeuN (red) in S1 of somatosensory cortex of male and female Cre (-) and Cre (+) mice. *Pten* KO neurons are mainly detected in L5. Scale bars=250 µm **B.** Group data of % of PTEN+ L5 neurons in somatosensory cortex of female and male Cre (+) mice as compared to same sex Cre (-) mice. n=7-14 sections from 3 mice/sex/genotype. **C.** Representative western blots for PTEN in microdissected L5 from somatosensory cortex from males and females Cre (-) and Cre (+) mice. GAPDH is used as a loading control. **D.** Quantification of PTEN levels in L5 of Cre(+) male and female mice, normalized to same sex Cre (-) littermates. (Females; n = 10 Cre(+)/Cre(-) pairs; Males; n=6 pairs).

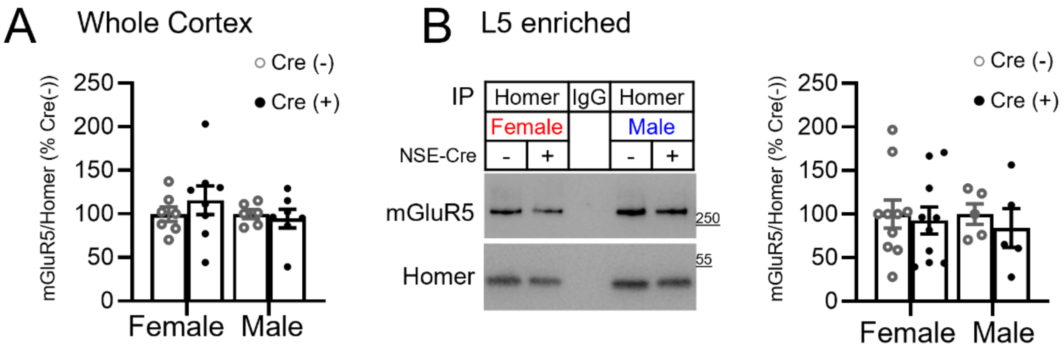


**Supplemental Figure 2. mGluR5-Homer interactions are unchanged in $^{NSE}$*Pten* KO mice. A.** Quantified group data of mGluR5-Homer co-immunoprecipitation (co-IP) from whole cortical lysates from female and male Cre (+) mice. **B.** Western blots of mGluR5-Homer co-IP from L5 microdissected (enriched) cortical lysates from male and female Cre(-) and Cre(+) mice. C. Group data from mGluR5-Homer co-IP from L5 cortical lysates.

Supplemental Figure 3

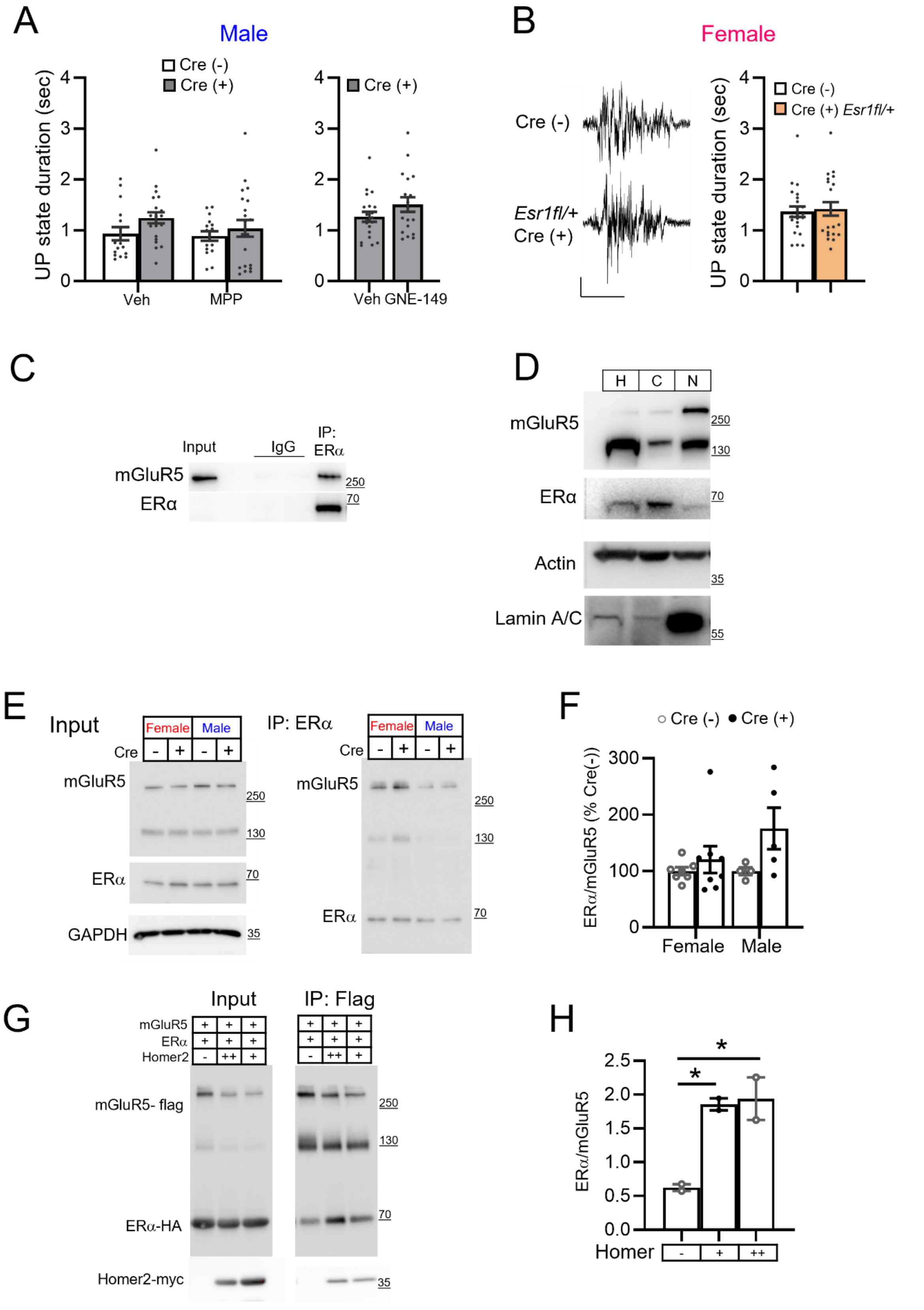

**Supplemental Figure 3. Genetic reduction of ERα in L5 neurons does not affect UP state duration in mice without *Pten* deletion, subcellular fractionation of mGluR5 and ERα and co-IP of mGluR5-ERα in $^{NSE}$*Pten* KO mice. A.** Group data show that the two ERα selective antagonist (MPP and GNE 1 μM; 1.5-2 hours) do not reduce UP state duration in slices from Cre (+) male mice. N=16-21 slices from 3-4 mice/genotype. **B.** Left: Representative UP states from female Cre (-) mice and mice with genetic reduction of ERα in L5 neurons (Cre (+) *Esr1*$^{fl/+}$). Scale bars= 0.05 mV/1 sec**.** Right: Quantified group data show that genetic reduction of ERα expression has no effect on UP state duration in females without *Pten* deletion. N= 21-22 slices from 3 mice/genotype. **C.** Representative immunoblot showing a specific co-IP of ERα and mGluR5 from cortical lysates with the ERα antibody and a rabbit IgG isotype. **D.** Western blot of subcellular fractionation of P21 mouse cortex (H, whole cell homogenate) shows that mGluR5 can be detected in the nuclear fraction (N), but not ERα, which is mainly expressed in the membrane and cytosolic (C) fraction. Lamin A/C was used as a control for nuclear fraction enrichment. **E.** Western blots mGluR5-ERα co-IP from whole cortical lysates from female and male Cre (+) and Cre (-) mice. **F.** Group data reveal no effect of sex or genotype on mGluR5- ERα co-IP. N= 4-8 mice/sex/genotype. **G.** Western blots and **H.** quantification of group data show that co-expression of Homer2 in HEK-292T cells increases the co-IP of ERα and mGluR5; n = 2 cultures/condition. Increasing amounts of Homer2 plasmid were transfected [1µg, (+); 2 µg, (++)].

Supplemental Figure 4

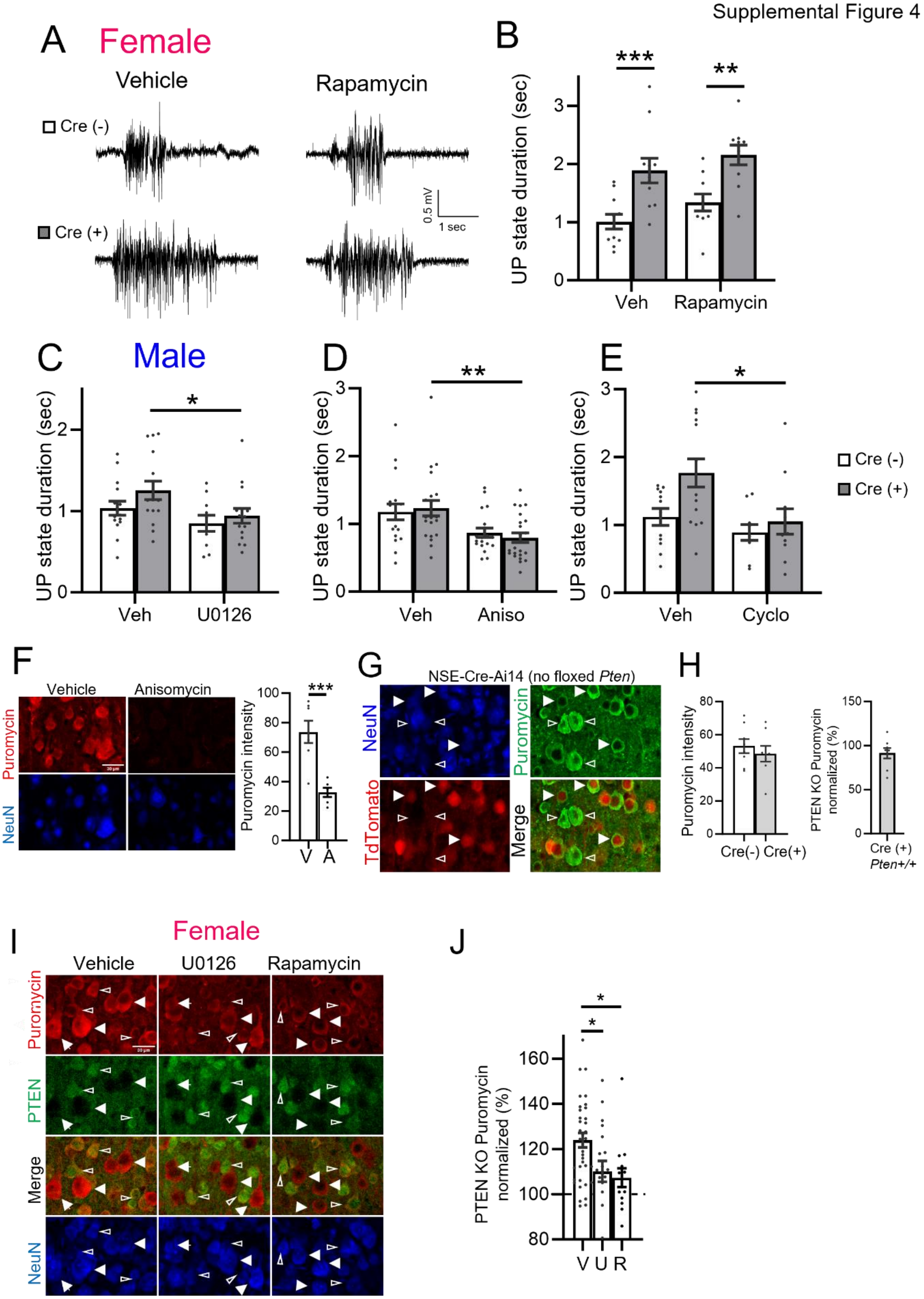

**Supplemental Figure 4. Effects of MEK/ERK and protein synthesis inhibitors on UP states in male $^{NSE}$*Pten* KO mice and puromycin labeling controls. A.** Representative UP states from female Cre (-) and Cre(+) mice pre-incubated (2 hr) in vehicle or rapamycin (200 nM). Scale bar = 0.5mV/1 sec. **B**. Group data of average UP state duration show that rapamycin pretreatment does not correct prolonged UP states in slices from female Cre (+) mice and has no effect on UP state duration in Cre (-) mice. N= 10-11 slices from 3 mice/genotype/condition. **C, D, E**. Group data of average UP state duration show that inhibitors of ERK activation or protein synthesis reduce UP state duration in slices from male Cre (+) mice. N= 10-22 slices from 3-6 mice/genotype/condition. **F.** Puromycin immunolabeling in neurons was reduced by pretreatment with the protein synthesis inhibitor, anisomycin. Representative images of NeuN and puromycin labeling and quantification of puromycin incorporation. Scale bar: 30 µm. N= 6-7 sections/condition from 1 mouse. Unpaired t-test. **G.** Representative images of puromycin immunolabeling (green) of L5 neurons in cortical sections from female NSE-Cre (+) mice with no *Pten*$^{fl/fl}$. NSE-Cre mice were crossed to a TdTomato Cre reporter line (Ai14) and endogenous TdTomato was used to identify Cre+ neurons (Filled arrows) and Cre(-) neurons (open arrows). Scale bar: 30 µm. **H.** Quantified group data show similar levels of puromycin immunolabeling in Cre(-) and Cre(+) cells without *Pten* deletion. N= 8 sections from 1 mouse. **I.** Representative images of puromycin immunolabeling (red) of L5 neurons in cortical sections from female Cre (+) mice pre-treated with vehicle or U0126 (20 µM; 2 hr) or rapamycin (200 nM; 2 hr). NeuN (blue) and PTEN (green) immunolabeling identify PTEN KO (filled arrows) and neighboring PTEN+ neurons (open arrows). Scale bar: 30 µm. **J.** Quantified group data of puromycin fluorescent intensity of PTEN KO neurons normalized to PTEN+ neurons within the same section. Puromycin intensity of PTEN KO neurons was similarly reduced by U0126 and rapamycin. N= 14-39 sections from 2-4 mice/treatment condition. One-way ANOVA and Sidak's multiple comparison test.

### Supplemental Figure 5

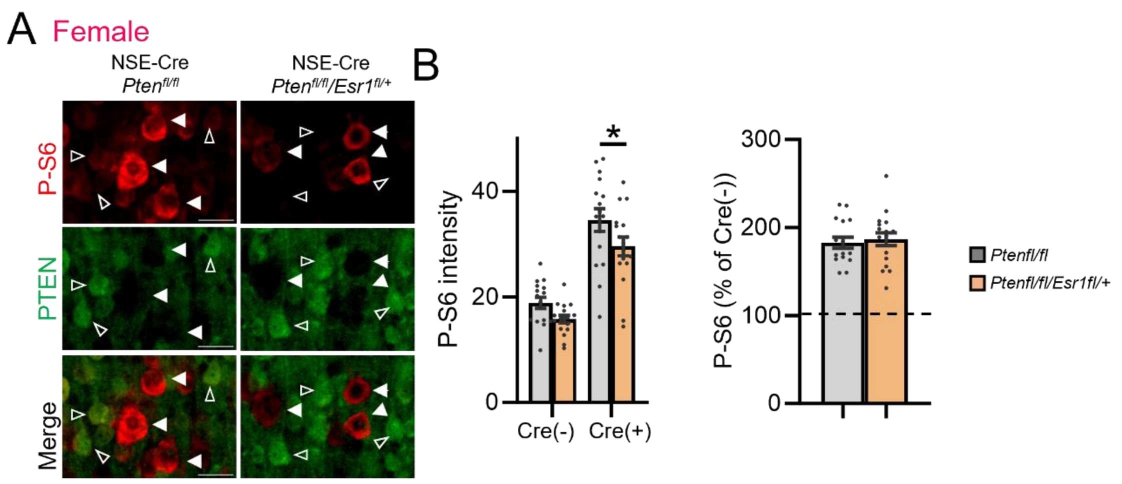


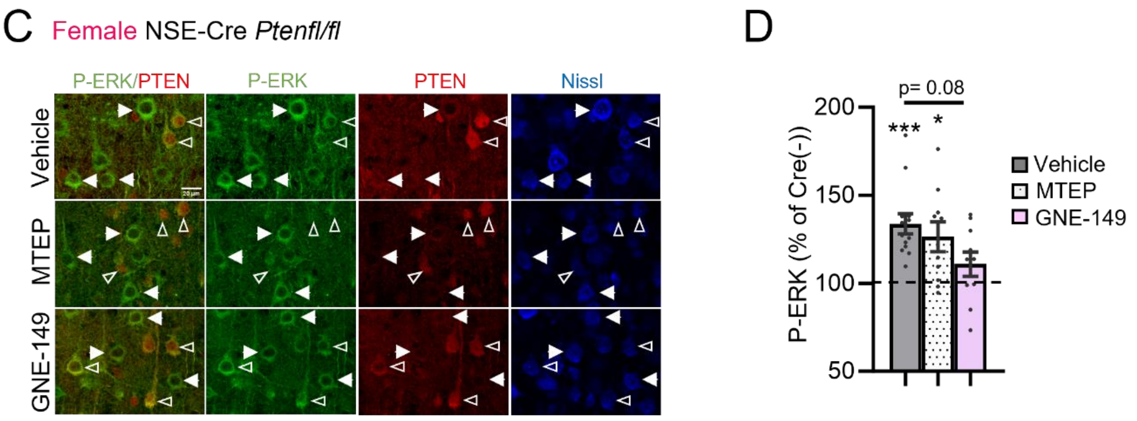


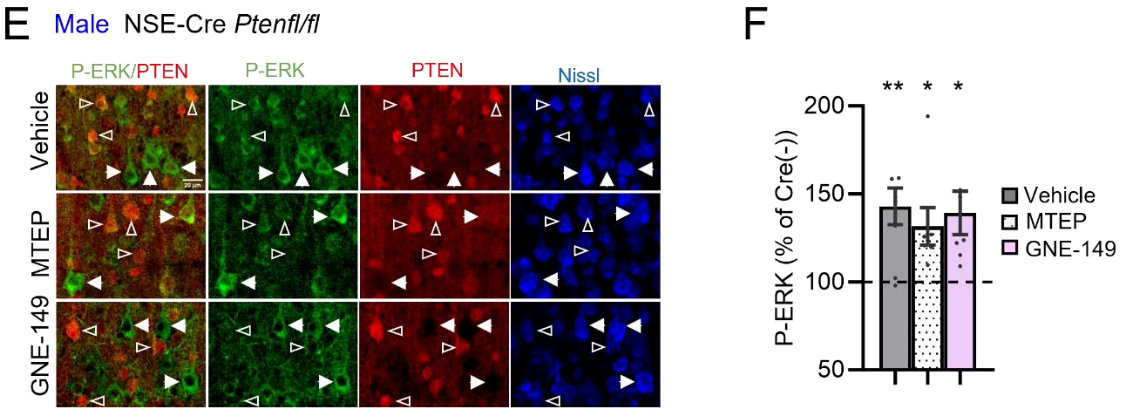

**Supplemental Figure 5. Heterozygous deletion of ERα reduces phosphorylated S6 in PTEN KO L5 neurons. A.** Representative images of phosphorylated (P) S6 (Ser235/236; red) and PTEN (green) in L5 cortical neurons from female NSE-Cre/*Pten*$^{fl/fl}$ or NSE-Cre/*Pten*$^{fl/fl}$/*Esr1*$^{fl/+}$ mice. Scale bar: 30 µm. **B.** Left: Raw P-S6 intensity values show an increase in P-S6 in PTEN KO (Cre(+)) neurons that are reduced by genetic reduction of *Esr1*. Right: Normalizing P-S6 intensities in PTEN KO neurons to neighboring PTEN (+), or WT, neurons reveal no effect of *Esr1* reduction on P-S6. N= 19-20 sections/4 mice/genotype. **C,E.** Representative images of P-ERK staining (green) of L5 neurons in cortical sections from female (C) and male (E) NSE-Cre/*Pten*$^{fl/fl}$ mice. PTEN (red) immunolabeling identify PTEN KO (filled arrows) and neighboring PTEN+ neurons (open arrows). Scale bar = 20 µm. **D, F.** Quantified group data of P-ERK fluorescent intensity of PTEN KO neurons (normalized to PTEN+ neurons within the same section) show that P-ERK is enhanced in both male and female PTEN KO neurons and blocking the activity of ERα only reduces this enhancement in female but not male mice. N = 7-13 sections from 3 mice/sex. Kruskal-Wallis with multiple comparison and One Sample Wilcoxon test.

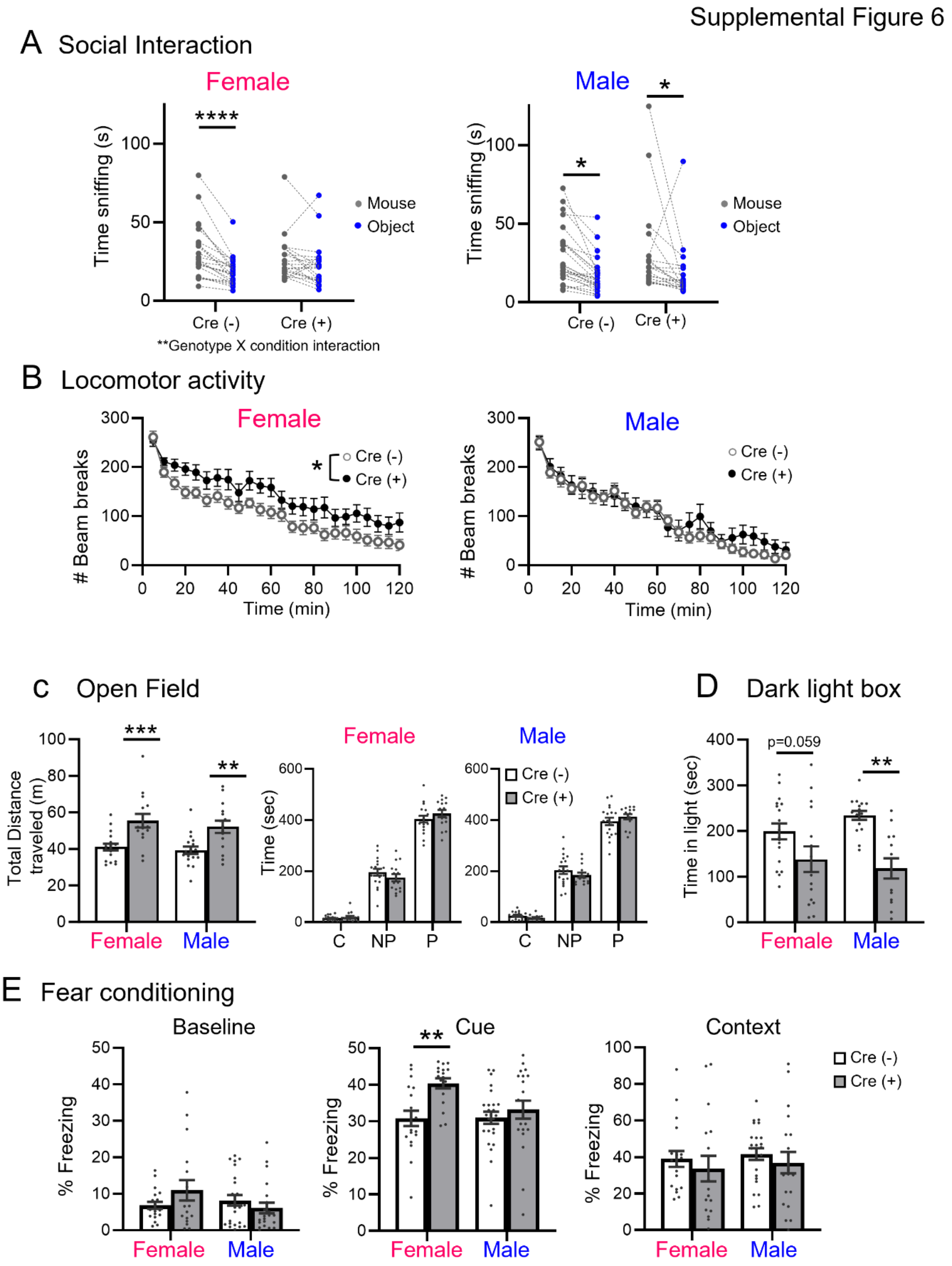


**Supplemental Figure 6. Female specific behavioral alterations in $^{NSE}$*Pten* KO mice. A.** Reduced sociability, measured as time sniffing, in the three-chamber social interaction test in female, but not male, Cre (+) mice. N= 20-26 mice/sex/genotype. **B.** Female specific increase in locomotor activity in Cre (+) mice measured as infrared beam breaks. (n= 15-18 mice/sex/genotype). **C.** Male and female Cre(+) mice were hyperlocomotive in an open field when assessed by total distance traveled. No sex or genotype differences were observed in time spent in the center (C), non-periphery (NP) or periphery (P) of open field. N=15-18 mice/sex/genotype. **D.** Cre(+) mice spent less time in the light side in the dark-light box test. N=13-18 mice/sex/genotype. **E.** Freezing to the cue in delay fear conditioning is enhanced in female, but not male, Cre (+) mice. Baseline and context-dependent freezing are not different between sex or genotype. N= 17-25 mice/sex/genotype.

Supplemental Figure 7

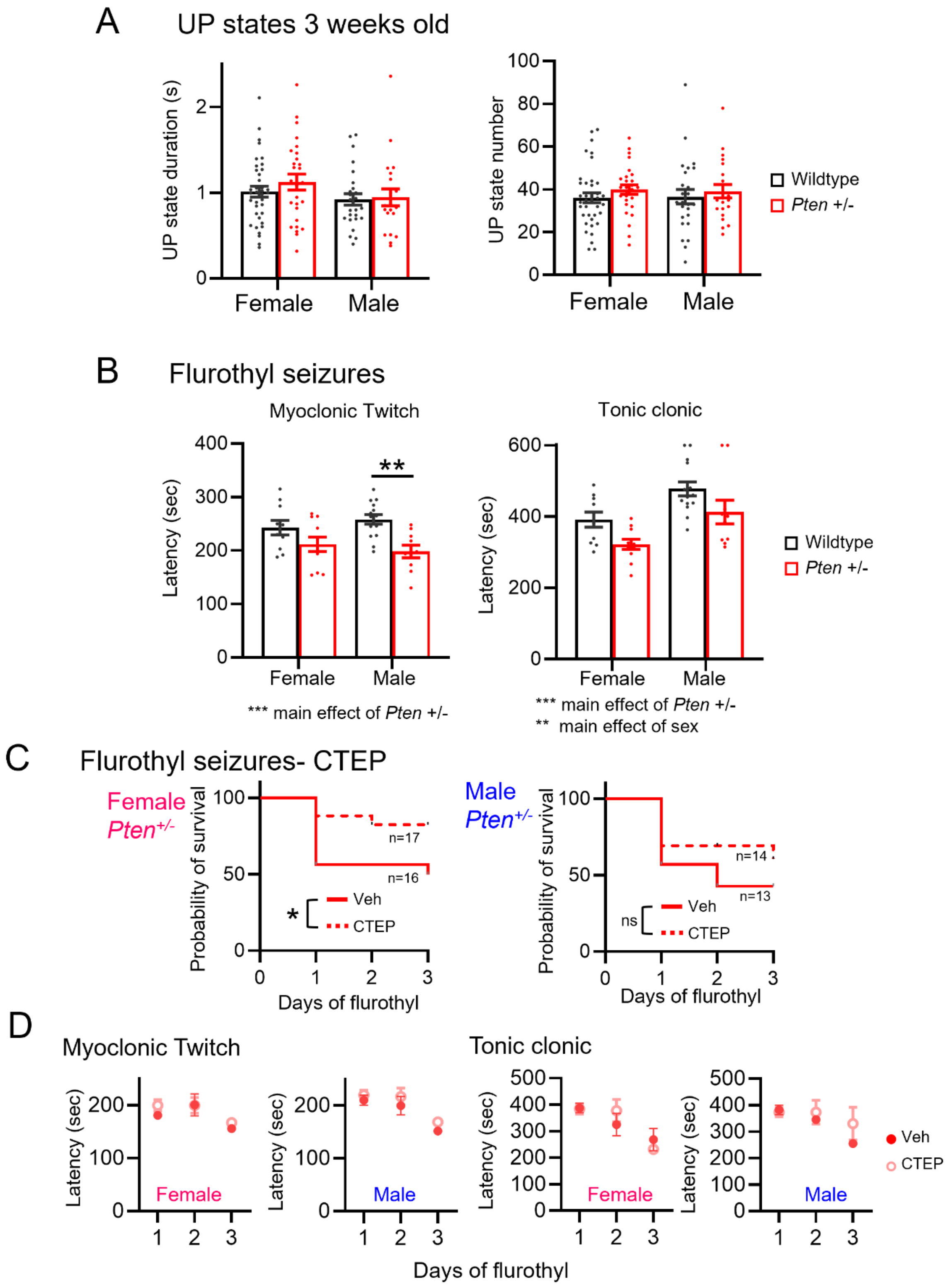

**Supplemental Figure 7. UP states in young *Pten*$^{+/-}$ mice and flurothyl seizure phenotypes in older *Pten*$^{+/-}$ mice. A.** Group averages reveal normal UP state duration (left) and number (in 5 min of recording) (right) in slices from P19-P22 female or male *Pten*$^{+/-}$ mice. **B.** Latency to myoclonus (left) and Tonic clonic seizure (right) in *Pten*$^{+/-}$ males and female mice. Main effects of sex and genotype are observed as indicated. **C**. Daily pretreatment with CTEP increases survival in female, but not male, *Pten*$^{+/-}$ mice across 3 days of flurothyl exposure. N=13-17 mice/sex/treatment condition. Gehan-Breslow-Wilcoxon test. **D.** Latency to myoclonic twitch and tonic-clonic seizure onset in *Pten*$^{+/-}$ mice are unaffected by pretreatment with CTEP.
